# *TSC1* loss-of-function increases risk for tauopathy by inducing tau acetylation and preventing autophagy-mediated tau clearance

**DOI:** 10.1101/2020.11.08.371922

**Authors:** Carolina Alquezar, Kathleen M Schoch, Ethan G Geier, Eliana Marisa Ramos, Aurora Scrivo, Kathy Li, Andrea R Argouarch, Elisabeth E Mlynarski, Beth Dombroski, Jennifer S Yokoyama, Ana M Cuervo, Alma L Burlingame, Gerard D Schellenberg, Timothy M Miller, Bruce L Miller, Aimee W Kao

## Abstract

Age-associated neurodegenerative disorders demonstrating tau-laden intracellular inclusions, including Alzheimer’s disease (AD), frontotemporal lobar degeneration (FTLD) and progressive supranuclear palsy (PSP), are collectively known as tauopathies. The vast majority of human tauopathies accumulate non-mutant tau rather than mutant forms of the protein, yet cell and animal models for non-mutant tauopathies are lacking. We previously linked a monoallelic mutation in the *TSC1* gene to tau accumulation and FTLD. Now, we have identified new variants in *TSC1* that predisposed to other tauopathies such as AD and PSP. These new *TSC1* risk variants significantly decreased the half-life of TSC1/hamartin *in vitro*. Cellular and murine models of *TSC1* haploinsufficiency (*TSC1*+/-) accumulated tau protein that exhibited aberrant acetylation on six lysine residues. Tau acetylation hindered its lysosomal degradation via chaperone-mediated autophagy leading to neuronal tau accumulation. Enhanced tau acetylation in *TSC1*+/- models was achieved through both an increase in p300 acetyltransferase activity and a decrease in SIRT1 deacetylase levels. Pharmacological modulation of either enzyme restored tau levels. Together, these studies substantiate *TSC1* as a novel tauopathy risk gene and advance *TSC1* haploinsufficiency as a new genetic model for tauopathy. In addition, these results promote acetylated tau as a rational target for diagnostic and therapeutic modalities in multiple tauopathies.

## Introduction

Tauopathies constitute a group of age-related neurodegenerative diseases characterized by the presence of neuronal and/or glial inclusions composed of the microtubule-binding protein tau. This group encompasses primary tauopathies, such as frontotemporal dementia FTD, corticobasal degeneration (CBD) and progressive supranuclear palsy (PSP) where tau inclusions are the major neuropathologic abnormality, as well as secondary tauopathies, such as Alzheimer’s disease (AD), where tau deposits occur in association with other pathologies (*1*). The incidence of tau-related neurodegenerative diseases is increasing, partly owing to the rise in life expectancy, but effective therapies are still lacking. Cumulative evidence from cellular and murine tauopathy models suggests that with age, the clearance of tau becomes impaired (*2–5*). However, many of these studies were performed in models expressing mutant, truncated or overexpressed forms of tau, while the majority of human tauopathies accumulate non-mutant forms of tau. Thus, unveiling the etiology of tauopathies requires models of endogenous nonmutant tau accumulation. Indeed, the limited understanding of the molecular basis for nonmutant tau accumulation has surfaced as a key impediment to the development of diagnostics and therapies for age-associated tauopathies.

Over the last two decades, an interesting pattern has emerged in which dose-dependent mutations in the same gene can predispose to both neurodevelopmental and age-associated neurodegenerative disorders. The first gene to be recognized as such was *glucocerebrosidase A* (*GBA*), in which mutations can underlie both Gaucher’s and Parkinson’s disease (*6*), but many others have since been identified (*7–10*). Recently, we described an individual with FTD who carried a novel, monoallelic loss-of-function (LOF) mutation in the *TSC1* gene (*11*). Until this report, *TSC1* mutations had only been associated with tuberous sclerosis complex (TSC), a juvenile-onset neurodevelopmental disorder typified by seizures, cognitive delay, space-occupying tumors and skin stigmata (*12*). As an adult-onset tauopathy gene, *TSC1* mutations would be another gene linking disorders widely separated by age. TSC can manifest in a highly heterogeneous manner, ranging from cases characterized by early childhood onset of intractable seizures and severe developmental delay to those presenting in adulthood with only skin stigmata and mild psychiatric symptoms (*13, 14*). To determine if otherwise cognitively normal adults with TSC exhibited any evidence for pre-morbid cognitive decline, we gathered a cohort of adults with mild TSC and found an age-associated cognitive phenotype consistent with FTD (*15*). In our earlier studies, *TSC1* mutations carriers exhibited evidence of tau accumulation in cell-based models, neuropathology (*11*) or by positivity on tau positron emission tomography (PET) imaging (*15*). Thus, clinical, genetic, experimental and neuroimaging data all support an association of *TSC1* gene mutations with tauopathy. However, additional genetic and mechanistic evidence would solidify *TSC1* this association.

The major objectives of this study were to validate *TSC1* gene as a tauopathy risk-factor and to unravel the mechanistic basis for tau accumulation associated with *TSC1* mutations. To do so, we leveraged the genetics of human tauopathy cohorts and identified additional *TSC1* risk variants that shorten the half-life of the TSC1/hamartin protein. We also generated murine and neuronal models of *TSC1* haploinsufficiency (*TSC1*+/-) and showed that they accumulated abnormally acetylated tau protein. Acetylation of tau prevented its efficient degradation in lysosomes. *TSC1* haploinsufficiency both promoted p300 histone acetyltransferase (HAT) activity and dampened SIRT1 deacetylase (HDAC) levels. Reversing these effects prevented tau accumulation. Collectively, this study reveals *TSC1* as a novel risk gene for tauopathies and identifies the mechanisms by which TSC1/hamartin haploinsufficiency leads to tau accumulation, which can be pursued in future pre-clinical studies focused on the development of effective treatments for tauopathies.

## Results

### Genetic variants in the TSC1 gene are overrepresented in cohorts of sporadic tauopathy

We have previously demonstrated a link between FTLD and pathogenic variants in *TSC1* (*11, 15*). To better understand the relationship between *TSC1* and risk of tauopathies, we examined the frequency of rare *TSC1* variants in a cohort of individuals diagnosed with early-onset Alzheimer’s disease (EOAD) (*16*), which has a stronger contribution of tau pathology than its more common, late-onset counterpart. The EOAD cohort showed an enrichment of the rare variant in *TSC1* rs2234980 dupTGC (also known as rs118203743), relative to controls from the Genome Aggregation Database (gnomAD v2.1.1) (minor allele frequency, MAF EOAD = 0.0066 vs MAF gnomAD = 0.00006). This variant results in a serine duplication at position 1043 (S1043dup) of the TSC1/hamartin protein. In light of this finding, we screened for *TSC1* variants in a cohort of pathologically confirmed PSP/tauopathy cases, compared with a control cohort from the Alzheimer’s Disease Sequencing Project (ADSP) Discovery dataset. The *TSC1* S1043 duplication (rs2234980) variant was also enriched in PSP cases, although not significantly (p-value = 0.4618, Odds Ratio = 1.897) **(Fig. 1A)**. In addition, another rare-coding variant in the *TSC1* gene (rs118203742), resulting in a glycine to serine change at position 1035 of the protein (G1035S), was found to be significantly associated with PSP (p-Value = 0.031, Odds Ratio = 4.241) (**Fig. 1A**). Sanger sequencing analysis confirmed the presence of the variants and showed that both were heterozygous. These results demonstrate that *TSC1* gene may be associated with increased risk for tauopathies.

**Figure 1.**
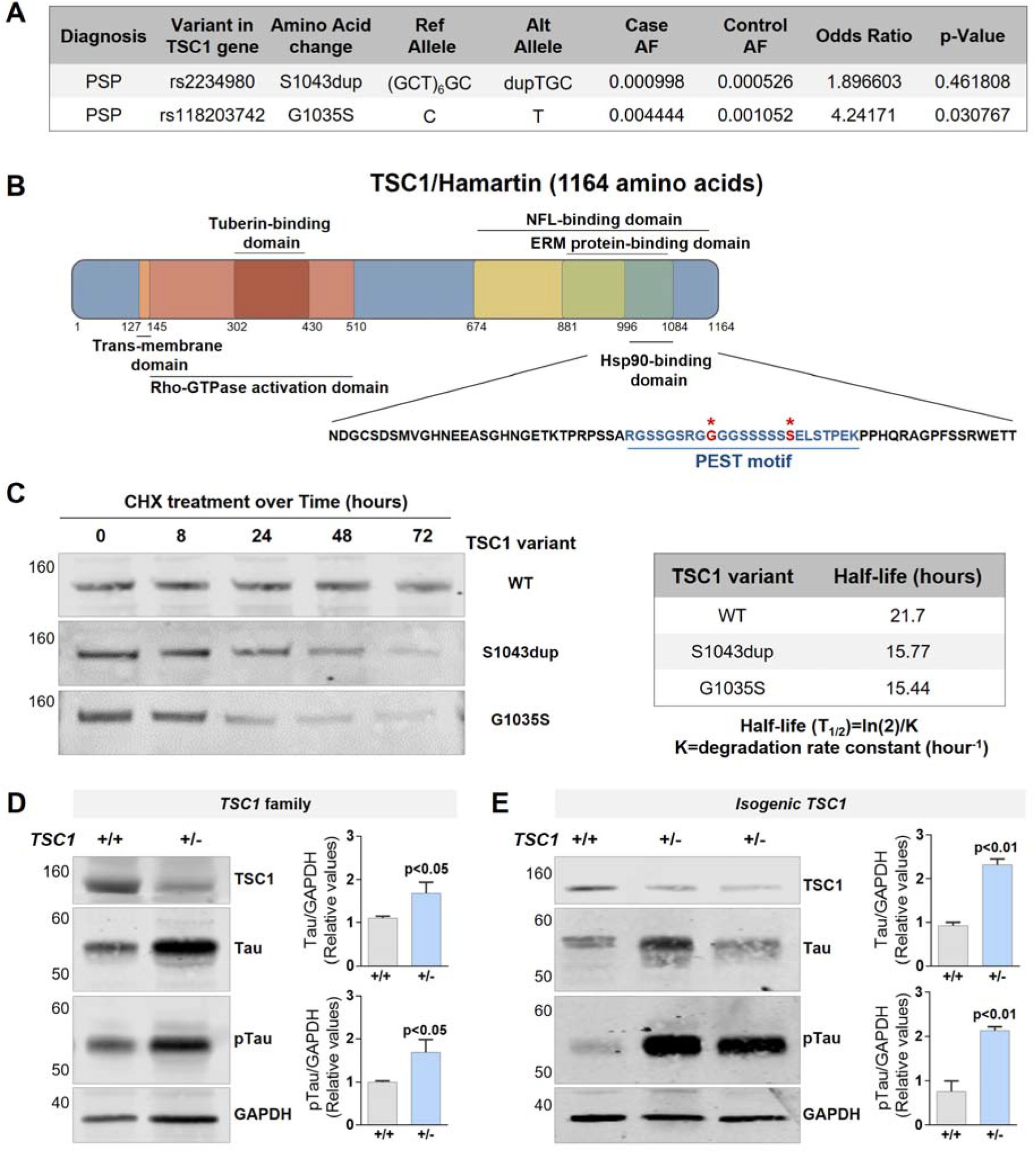
Genetic variants in *TSC1* are overrepresented in tauopathy patients. **A**) Allelic frequency, Odds ratio and p-value of the *TSC1* genetic variants found in the pathologically confirmed PSP cases compared to healthy control individuals from the ADSP database. **B**) Schematic representation of TSC1/hamartin protein with functional domains identified. The Hsp90-binding domain sequence is indicated with PEST motif in dark blue and the position of the disease-associated variants marked with a red asterisk. **C**) Stable SH-SY5Y cells lines expressing FLAG-tagged WT or mutant TSC1/hamartin protein were treated with 50μg/mL of CHX, and lysates were collected at different time points. A”ti-FLAG immunoblots show the levels of TSC1/hamartin before and after CHX treatment. TSC1/hamartin half-life is shown in the table to the right. **D-E**) Immunoblots showing the levels of TSC1/hamartin protein, total tau and phosphorylated tau (S396/S404) in iNeurons differentiated from iPSC from a family of *TSC1* mutation carriers (D) and *TSC1*+/- isogenic iPSCs generated in the background of the PGP1 line (E). Plots show the mean ± SEM of three independent experiments carried out with all the cell lines listed in Table S2. Statistical significance was assessed by Student’s t-test. Differences were considered statistically significant when p<0.05.

The *TSC1* gene encodes for a large, 1164 amino acid protein known as TSC1/hamartin. Interestingly, both the rs2234980 and rs118203742 variants alter the coding sequence of the region of TSC1/hamartin implicated in binding to the Hsp90 chaperone protein (*17*). Within this Hsp90-binding region, we further identified a potential PEST motif (<u>ePESTfind tool</u>) (*18*) (**Fig. 1B**). PEST motifs are amino acid sequences enriched with proline (P), glutamate (E), serine (S) and threonine (T) residues that regulate protein degradation via the proteasome (*18*). Given their localization within the PEST motif, we posited that the tauopathy-associated *TSC1* variants could exert a loss-of-function effect by disrupting the PEST motif, thereby shortening the half-life of TSC1/hamartin. To test this possibility, we generated cell lines expressing tagged versions of wild-type (WT) or mutant TSC1/hamartin. Cells were treated with cycloheximide (CHX) to halt protein synthesis and the half-life of WT and mutant TSC1/hamartin protein was measured over time. Both of the tauopathy-associated *TSC1* genetic variants shortened TSC1/hamartin protein half-life by nearly 30% (**Fig. 1C**). These results suggest that, similar to our earlier reports (*11, 15*), genetic variants that decrease levels of TSC1/hamartin are associated with tauopathy.

### iNeuron models of TSC1 haploinsufficiency accumulate tau

We have previously shown that *TSC1* haploinsufficiency results in the specific accumulation of total and phospho-tau in the SH-SY5Y neuronal-like cell line (*11*). However, it remained unknown if *TSC1* haploinsufficiency also induces tau accumulation in iPSC-differentiated iNeurons. To assess this, we generated induced pluripotent stem cells (iPSCs) carrying *TSC1*+/- mutations. We obtained fibroblasts from three members of a family carrying a heterozygous LOF mutation in *TSC1* associated with FTLD-tau as well as one WT control family member (*11*) (**Fig. S1**). The fibroblasts were reprogrammed into iPSCs, differentiated into iNeurons and compared to a well-characterized WT iPSC line (F11350) (*19, 20*) (**Table S2**). As expected, TSC1/hamartin levels were decreased in the *TSC1*+/- lines compared to WT controls. In addition, *TSC1*+/- iNeurons exhibited increased levels of total tau and phospho-tau compared to controls (**Fig. 1D**).

Although the patient-derived cell lines are a useful proof of concept that humanized neuronal models of *TSC1* haploinsufficiency also accumulate tau, these lines suffer from the limitation of differences in genetic background. Thus, we also utilized CRISPR/Cas9 genome editing to generate *TSC1*+/- lines in the background of the PGP1 WT parent iPSC line (**Fig. S2** and **Table S2**). The resulting iPSC lines were isogenic at all but the *TSC1* locus. Two independent *TSC1*+/- clones were generated, differentiated into iNeurons and used to assess tau compared to the parent line. Similar to the *TSC1* family cell lines, these isogenic *TSC1*+/- neurons demonstrated increased total tau and phospho-tau levels (**Fig. 1E**). Thus, *TSC1* haploinsufficiency induces tau accumulation in both patient-derived and engineered cell lines compared with isogenic controls, supporting the results previously found in TSC1+/- neuronal-like SH-SY5Y cells.

### Tau accumulation in TSC1 haploinsufficiency is not a consequence of impaired autophagy activity

One of the functions of TSC1/hamartin protein is to stabilize TSC2/tuberin (*21*), a GTPase-activating protein that negatively regulates the mTOR (mammalian target of rapamycin) pathway (*21*). Therefore, TSC1/hamartin haploinsufficiency leads to TSC2/tuberin degradation and mTOR overactivation (**Fig. S3A**). The mTOR pathway controls important cellular functions including protein degradation through autophagy (*22*), and thus, overactivation of mTOR downregulates autophagy and impairs protein degradation (**Fig. S3A**). Several studies have reported that mTOR activation is associated with neurodegenerative diseases (*23, 24*), and that tau protein is degraded in lysosomes via autophagy (*25, 26*). In this context, we wondered if tau accumulation in TSC1/hamartin haploinsufficiency was due to an overall decrease in protein degradation through the autophagy pathways (autophagic flux) due to abnormal mTOR activation.

To evaluate this possibility, we first considered macroautophagy, the best characterized type of autophagy, and measured autophagic flux in a series of fibroblasts with and without a LOF mutation in *TSC1* (*11*) (**Fig. S3B-C**). Cells were transduced with lentivirus carrying an mCherry-GFP-LC3 tandem construct that allows measuring flux as the maturation of autophagosomes (vesicles positive for mCherry and GFP fluorescence) into autolysosomes (vesicles in which GFP fluorescence is loss because of acidification upon fusion with lysosomes) (*27*). Fibroblasts carrying the *TSC1* mutation (*TSC1*+/-) demonstrated higher abundance of autophagic vacuoles (overall autophagic compartments independently of their maturation state). This increase was driven, for the most part, by autolysosomes, supporting an increase, rather than a decrease, in macroautophagic flux associated with TSC1/hamartin haploinsufficiency (**Fig. 2A**).

**Figure 2.**
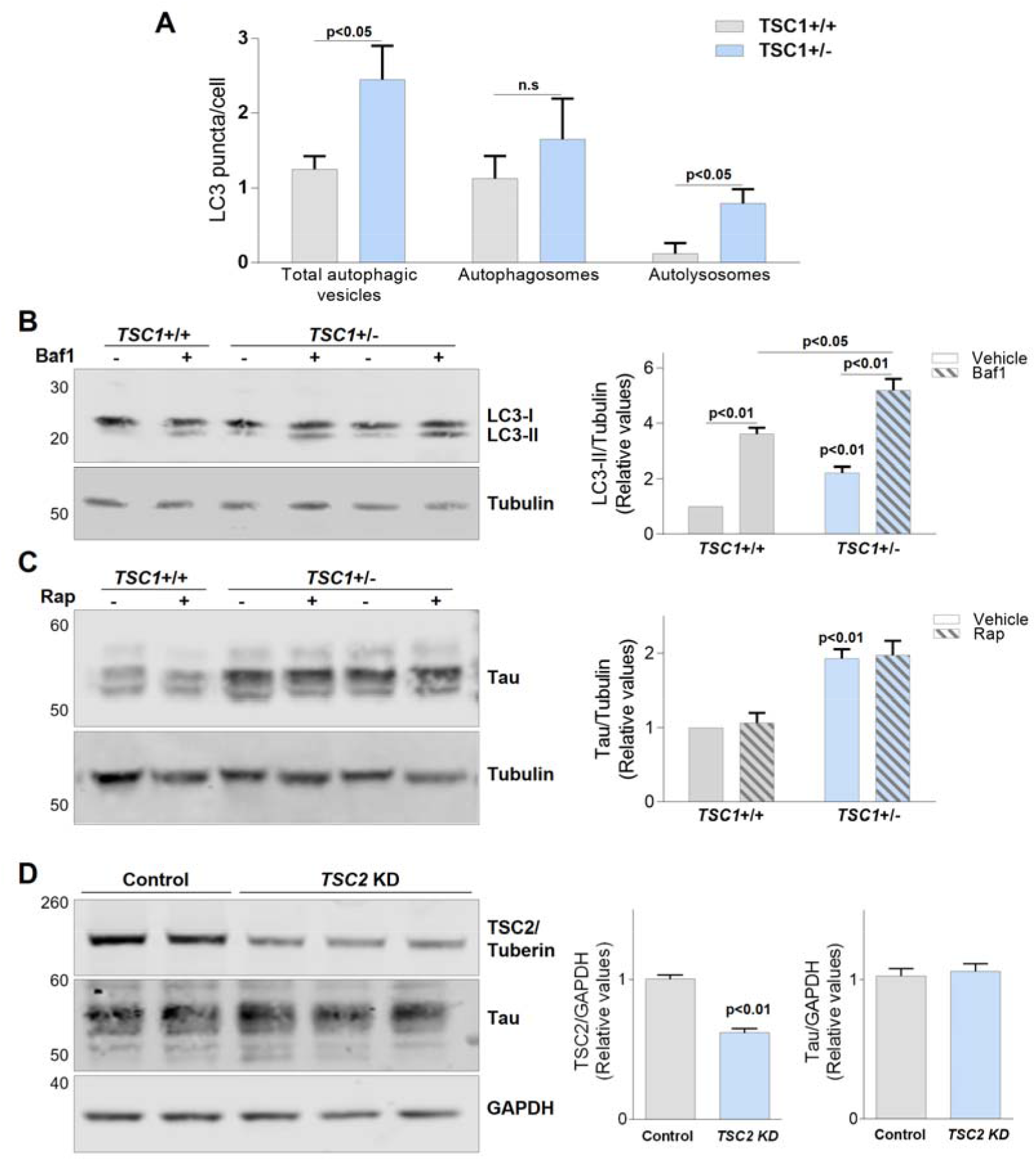
TSC1/hamartin haploinsufficiency induces tau accumulation independent of macroautophagy impairment and mTOR pathway activation. **A**) Plot showing the number of LC3-II puncta per cell in *TSC1*+/+ and *TSC1*+/- fibroblasts. Total autophagic vesicles consist of autophagosomes and autolysosomes. **B**) Anti-LC3 immunoblot showing LC3-I and II levels before and after 6 hours of treatment with 100nM of bafilomycin 1 (Baf1) in control or *TSC1*+/- differentiated SH-SY5Y cells. **C)** Immunoblot showing tau levels before and after 72 hours of rapamycin (Rap, 25nM) treatment in WT or *TSC1*+/- differentiated SH-SY5Y cells. **D)** Representative immunoblots showing the levels of TSC2/tuberin and tau proteins in *TSC2* knockdown (KD) SH-SY5Y stable lines compared to controls. All plots show the mean ± SEM from three independent experiments. Two-way ANOVA tests followed by Bonferroni’s comparison (A-C) and one-way ANOVA test (D) were used to assess statistical significance. Differences were considered statistically significant when p<0.05. Unless otherwise indicated, p-values indicate comparison to the first, left-most column in the graph.

Overactivation of autophagy has been previously reported in early stages of AD (*28*), where it has been proposed to have a compensatory, neuroprotective role. To address whether *TSC1* haploinsufficiency caused a compensatory increase in autophagy in neuronal cells, we measured the levels of LC3-II protein in differentiated SH-SY5Y cells before and after treatment with bafilomycin (Baf1), a V-ATPase inhibitor that blocks protein degradation in the lysosomes. This approach provides an estimate of protein degradation through the autophagy-lysosome system (*29*). *TSC1*+/- SH-SY5Y cells exhibited increased LC3-II in the basal state, which further increased after Baf1 treatment (**Fig. 2B**). These results confirmed that in both fibroblasts and differentiated SH-SY5Y cells, *TSC1* haploinsuffíciency caused a compensatory enhancement in autophagic flux, rather than inhibition, as would be expected as result of the aberrant mTOR activation previously demonstrated (*11*).

We next asked whether increased mTOR activity is required for tau accumulation in *TSC1*+/- cells. First, control and *TSC1*+/- cells were treated with rapamycin, a specific mTOR inhibitor. Rapamycin treatment did not prevent tau accumulation in *TSC1*+/- cells (**Fig. 2C**). In addition, since TSC1/hamartin regulates mTOR activity through the stabilization of TSC2/tuberin, we generated TSC2/tuberin knockdown (KD) SH-SY5Y cell lines. Differentiated *TSC2* KD cells exhibited elevated mTOR activity, reflected by the increase in the phosphorylation status of P70S6k protein (**Fig. S3D**). However, *TSC2* KD did not increase the levels of total tau protein compared to control cell lines (**Fig. 2D**). Therefore, mTOR activation was neither necessary nor sufficient to induce tau accumulation in TSC1/hamartin haploinsufficient cells.

### TSC1/hamartin haploinsufficiency prevents tau degradation by chaperone mediated autophagy

Since tau accumulation in *TSC1*+/- cells was not due to a general decrease in autophagy, we next investigated how TSC1/hamartin specifically regulates tau levels. In general, protein levels can be regulated via production or degradation. Therefore, we first determined if tau expression changes in *TSC1*+/- cells. Tau (*MAPT*) mRNA levels were assessed by quantitative PCR in differentiated SH-SY5Y cells. *TSC1*+/- cells showed no difference in tau mRNA expression compared to controls (**Fig. 3A**).

**Figure 3.**
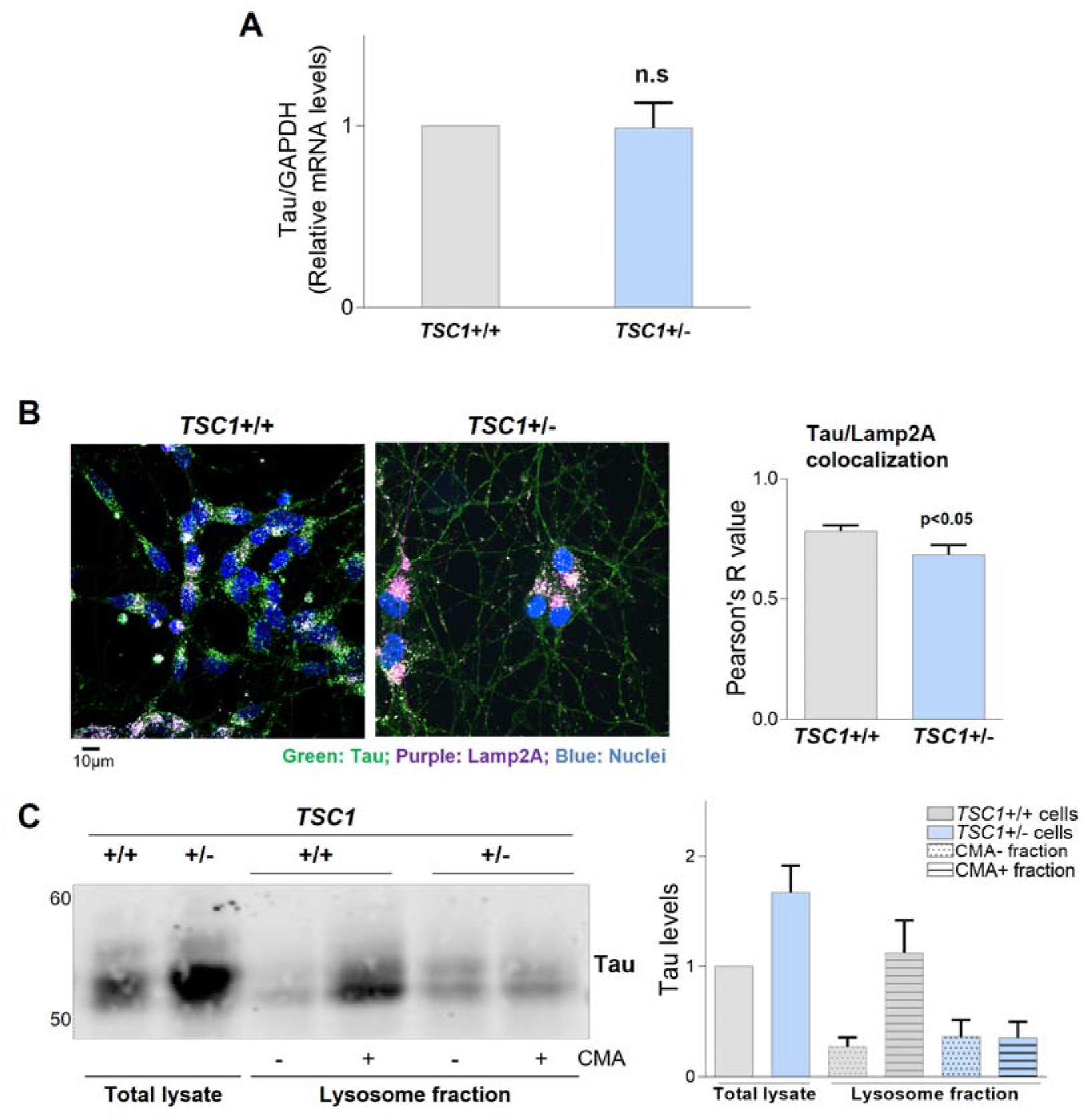
Loss of TSC1/hamartin impairs tau degradation in the lysosomes by chaperone mediated autophagy. **A**) Tau mRNA levels in WT and *TSC1*+/- differentiated SH-SY5Y cells measured by qPCR. **B**) Control and *TSC1*+/- differentiated SH-SY5Y cells were immunostained with antibodies against total tau (green) and the lysosomal marker Lamp2A (purple). Nuclei were stained with DAPI (blue). Ten different areas for each cell line were imaged, and the experiment was performed three times. Plot shows the average of Pearson’s R correlation value for both channels. The statistical significance of Pearson’s R correlation value was assessed by Student’s t-test. **C**) CMA active (+) and CMA inactive (-) lysosomal fractions were isolated from control and *TSC1*+/- differentiated SH-SY5Y cells. Representative immunoblot showing the distribution of tau protein in the lysosomal compartments is shown. Lysosomal isolation was performed twice with identical results. Plot shows the mean ± SEM of tau levels from two independent experiments. Differences were considered statistically significant when p<0.05.

We then turned our attention to tau clearance. Chaperone-mediated autophagy (CMA) is a form of selective protein degradation that requires recognition of a KFERQ motif by the Hsc70 chaperone and subsequent binding and translocation of the targeted protein into the lysosomes through the Lamp2A (lysosome-associated membrane protein type 2A) channel (*30–32*) Tau has two KFERQ motifs in its sequence, and thus is a client for CMA (*26*). To understand whether TSC1/hamartin haploinsuffíciency impairs tau degradation in lysosomes, control and *TSC1*+/- cells were immunostained with tau and Lamp2A antibodies and confocal microscopy was used to assess the colocalization. *TSC1*+/- cells showed decreased tau/Lamp2A colocalization compared with *TSC1+/+* cells (**Fig. 3B**), suggesting that TSC1 insufficiency impairs tau association with lysosomes. Since we did not find changes in tau degradation through macroautophagy, we propose that reduced tau association with lysosomes was mainly associated with CMA competent lysosomes. To directly test this, we performed biochemical separation of lysosome fractions CMA (*33*). CMA enriched (CMA+) and CMA non-enriched (CMA-) lysosomes were isolated from control and *TSC1*+/- cells and then were immunoblotted for tau. The TSC1/hamartin haploinsufficiency lead to a very pronounced decrease of tau levels in CMA+ lysosomes (**Fig. 3C** and **Fig. S4**). Thus, these results confirmed that in *TSC1*+/- cells, tau tau degradation via CMA+ is severely compromised.

### Tau acetylation impairs its degradation by chaperone mediated autophagy in TSC1+/- cells

Tau undergoes many posttranslational modifications (PTMs) that play an important role in its function and homeostasis (*34*). PTMs have the potential to directly regulate tau degradation via both the proteasome and the lysosomes (*35–37*). To explore whether TSC1/hamartin haploinsufficiency impacts tau PTMs, we performed tau immunoprecipitation (IP) followed by mass spectrometry analysis. Although phospho-tau accumulates in TSC1/hamartin haploinsufficiency, the actual residues that were phosphorylated on tau were not different in *TSC1*+/- cells compared to controls (**Fig. S5**). On the other hand, compared to WT cells, tau extracted from *TSC1*+/- cells was uniquely acetylated on six lysine (K) residues: K150, K240, K343, K369, K375 and K385 (**Fig. 4A-B** and **Fig. S5**). K174 was found to be acetylated in both control and *TSC1*+/- cells (**Fig. S5**).

**Figure 4.**
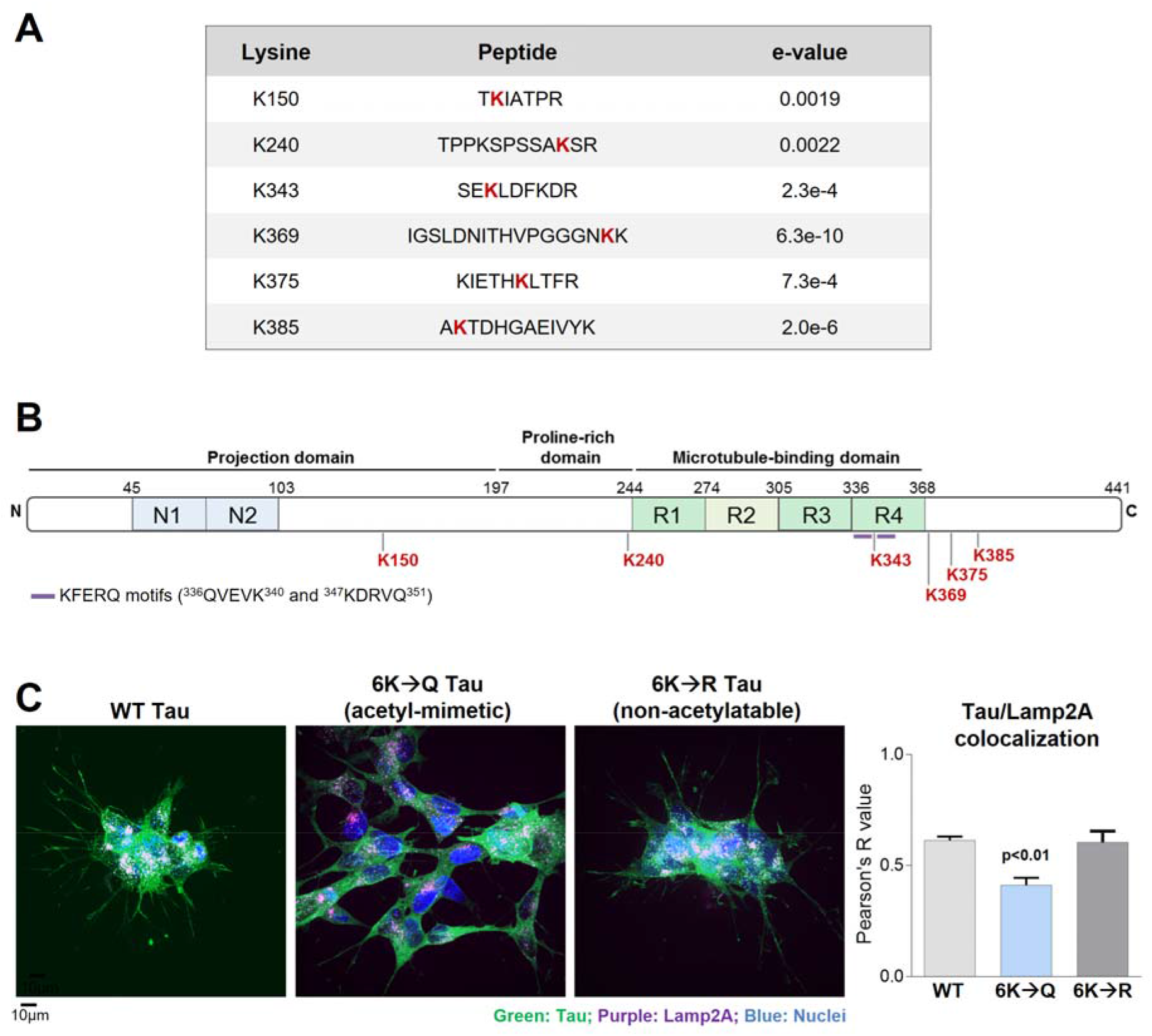
Tau acetylation in TSC1 haploinsufficient cells alters lysosomal tau association. **A**) List of acetylated lysine residues found by mass spectrometry in *TSC1*+/- but not WT SH-SY5Y cells. The acetylated lysine residue (K) is highlighted in red. The expectation value (evalue), a measure of the statistical significance of a peptide assignment, is presented in the right column of the table. **B**) Diagram of tau protein annotated with the most important functional domains, KFERQ motifs and the lysine residues presenting TSC1/hamartin-associated acetylation. **C**) SH-SY5Y cells expressing FLAG-tagged WT 2N4R tau, constitutive acetylated tau (2N4R 6K→Q tau) and non-acetylated tau (2N4R 6K→R tau) were stained with an anti-FLAG antibody recognizing overexpressed tau (green) and endogenous Lamp2 protein (purple). Nuclei were stained with DAPI (blue). Plot shows the average of Pearson’s R correlation value. Ten different areas for each cell line were imaged. Experiments were performed in triplicates. Statistical significance was assessed using one-way ANOVA. Differences were considered statistically significant when p<0.05.

Abnormal acetylation of mutant tau has been reported to interfere with its degradation and thus promote tau accumulation (*3, 38–41*). The acetyl-lysine residues identified in *TSC1*+/- cells have not previously been described on endogenous human non-mutant tau, and thus their functional impact on tau clearance is unknown. To determine if TSC1-related acetylation prevents tau degradation by CMA, we generated stable cell lines expressing 2N4R WT tau or 2N4R tau in which the six acetylated lysine residues in *TSC1*+/- cells were mutated to acetyl-mimetic glutamine residues (6K→Q Tau) or non-acetylatable arginine residues (6K→R Tau). If lysine acetylation prevented CMA-mediated degradation, one would predict that tau colocalization with Lamp2A would decrease. Indeed, tau/Lamp2A colocalization decreased in cells expressing 6K→Q acetyl-mimetic tau but not in cells expressing non-acetylatable 6K→R tau (**Fig. 4C**). These data demonstrate that the mechanistic basis for tau accumulation in TSC1/hamartin haploinsuffíciency is abnormal acetylation that subsequently disrupts tau degradation by CMA.

### TSC1/hamartin regulates p300 HAT and SIRT1 HDAC activities

Acetyl groups are conferred onto proteins by histone acetyltransferases (HATs) such as p300 and are removed by deacetylases (HDACs) such as SIRT1 (*3, 4, 42*) To better understand the process of tau acetylation in *TSC1* haploinsuffíciency, we tested p300 HAT activity and SIRT1 HDAC levels. *TSC1*+/- cells showed increased p300 HAT activity as measured by the acetylation of histone 3 on lysine 18 (H3^acK18^) (**Fig. 5A**). In addition, we also directly assessed HAT activity with a specific enzymatic assay that similarly showed increased p300 HAT activity in *TSC1*+/- cells (**Fig. 5B**). Next, we evaluated deacetylase activity. Accompanying the increased p300 acetylase activity, *TSC1*+/- cells also exhibited reduced levels of SIRT1 HDAC protein and mRNA (**Fig. 5C-D**). Thus, TSC1 haploinsuffíciency both enhances acetyltransferase and decreases deacetylase activity, which results in a net increase in tau acetylation.

**Figure 5.**
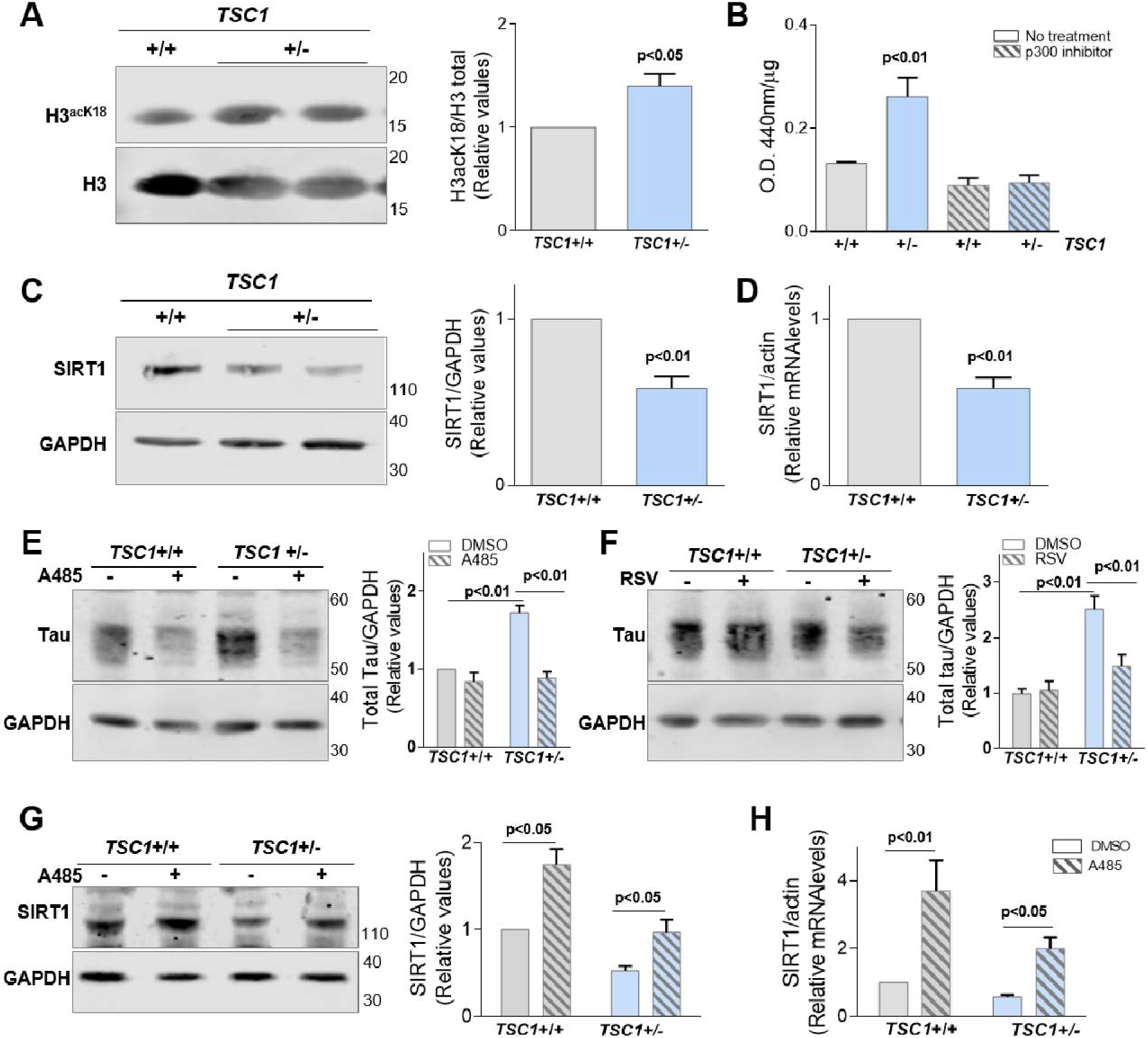
TSC1/hamartin insufficiency regulates p300 HAT activity and SIRT1 HDAC levels. **A**) p300 HAT activity in control or *TSC1*+/- differentiated SH-SY5Y cells indirectly assessed by the levels of histone 3 (H3) acetylated at lysine 18 (H3^acK18^) **B**) Enzymatic activity of HAT in differentiated SH-SY5Y cells before and after the treatment with a specific p300 inhibitor. **C-D**) SIRT1 protein (C) and mRNA levels (D) measured in control and *TSC1*+/- differentiated SH-SY5Y cells. **E-F**) Control and *TSC1*+/- differentiated cells were treated for 72 hours with a specific p300 inhibitor, A485 (15μM) (E) or with the SIRT1 activator, resveratrol (RSV) (25μM) (F). **G-H**) Effect of p300 inhibition on the levels of SIRT1 protein (G) and mRNA (H) in control and *TSC1*+/- differentiated SH-SY5Y cells. All experiments were performed in triplicates. Plots show the mean ± SEM for each experiment. Statistical significance was assessed using Student’s t-test (A-D) or two-way ANOVA (E-H) followed by Bonferroni’s comparison. Differences were considered statistically significant when p<0.05.

We next asked if modulation of p300 HAT and SIRT1 HDAC impacts tau accumulation in TSC1/hamartin haploinsuffíciency. First, we pharmacologically inhibited p300 HAT activity (with A485) (*43*) and found that it reduced tau levels in *TSC1*+/- cells to TSC1+/+ levels (**Fig. 5E**). We then activated SIRT1 deacetylase activity (with resveratrol, RSV) (*44*) and similarly found that it normalized tau levels in *TSC1*+/- cells (**Fig. 5F**). Tau mRNA expression was unaffected (**Fig. S6**). Interestingly, treatment with the p300 HAT inhibitor may decrease total tau both directly, via decreased p300 activity, and indirectly, by increasing the expression of the HDAC SIRT1 (**Fig. 5G-H**), implying that p300 HAT regulates SIRT1 expression. Taken together, these data demonstrate that TSC1/hamartin insufficiency results in increased p300 HAT and decreased SIRT1 HDAC activity and that correcting the dysregulation of these enzymes normalizes tau levels in *TSC1*+/- cells.

### TSC1/hamartin haploinsufficiency induces tau acetylation and neurodegeneration in vivo

To determine whether TSC1/hamartin impacts tau levels and post-translational modifications *in vivo* and in an age-dependent manner, we generated a model of neuronal TSC1/hamartin haploinsufficiency in mice expressing human tau (*hTau;TSC1*^Syn1+/-^). At 6 months, *hTau;TSC1*+/- animals were not significantly different from controls relative to tau pathology. However, at 15 months, *hTau;TSC1*+/- mice demonstrated a significant accumulation of phosphorylated tau protein in the retrosplenial cortex (RSCx), a region involved in hippocampal-cortex circuitry that functions in spatial cognition and memory (*45, 46*) and that is affected early in tauopathies such as AD (*47*) (**Fig. 6A**).

**Figure 6.**
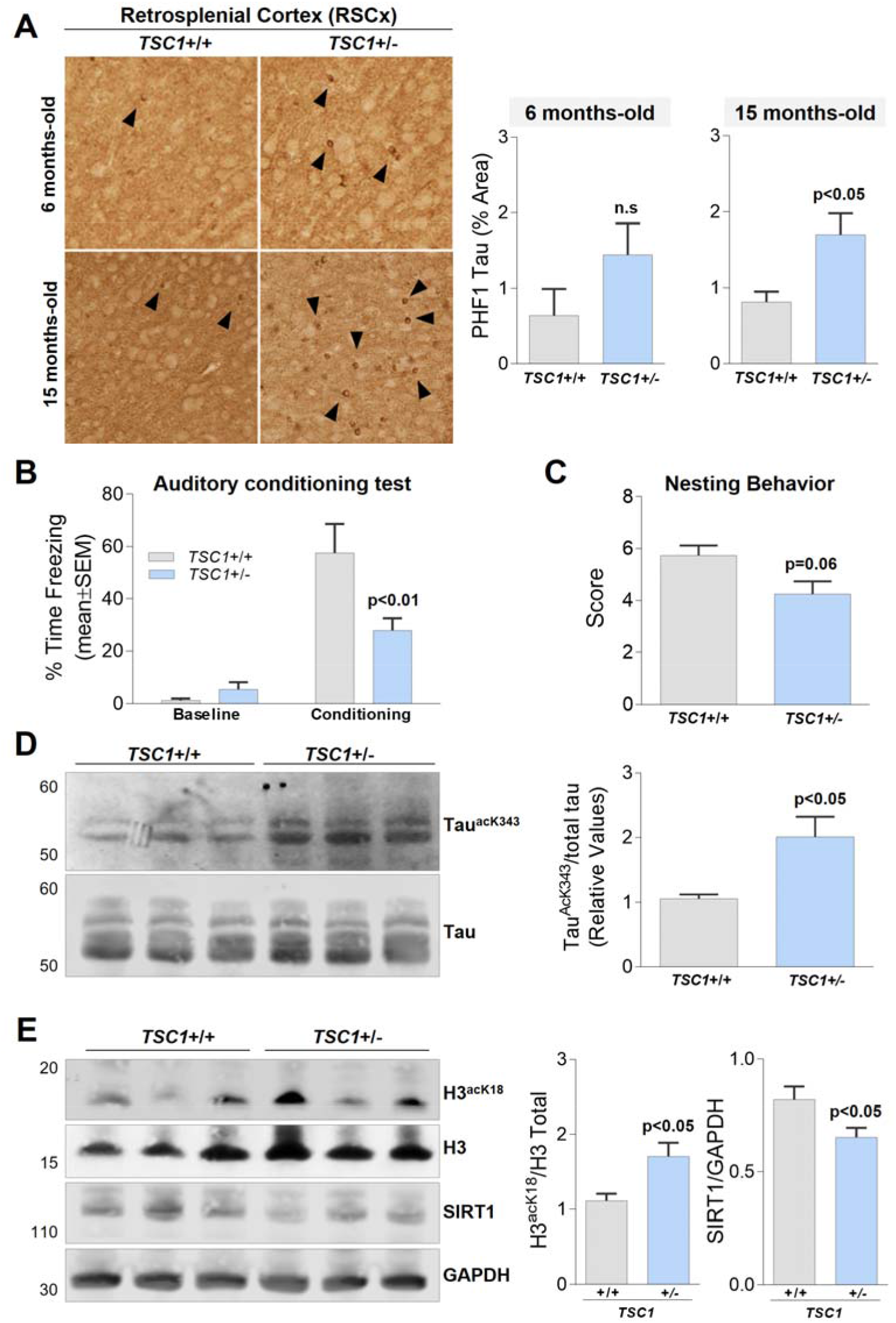
TSC1/hamartin insufficiency promotes tau acetylation and age-related neurodegeneration in mice. **A**) Tau immunostaining with PHF1 antibody recognizing tau phosphorylated at S396 and S404 residues was performed in the retrosplenial cortex (RSCx) of *TSC1*+/- (*hTau;TSC1*^Syn1+/-^) and control *TSC1*+/+ mice (*hTau;TSC1*^Syn1+/+^). Brains were stained at 6 and 15 months-old. Representative images of tau staining in RSCx are shown. Plots represent the percentage of the area of the brain positive for tau staining. Student’s t-test was used to assess statistical significance. **B**) Auditory conditioning test performed in 15 month-old mice at baseline and after nine minutes of conditioning. Two-way ANOVA followed by Bonferroni’s individual comparison was used to analyze statistical differences. **C**) Nesting behavior score of 15-month-old *TSC1*+/- (*hTau;TSC1*^Syn1+/-^) mice compared with control *TSC1*+/+ mice (*hTau;TSC1*^Syn1+/+^). Statistical analyses were performed using Student’s t-test. **D**) Immunoblots showing the levels of tau protein acetylated at lysine 343 (Tau^acK343^) and total tau in the RSCx of 15 month-old *TSC1*+/- mice. **E**) p300 HAT activity was measured in the RSCx of 15 month-old mice by assessing the acetylation status of H3 at K18. SIRT1 levels were measured in the RSCx of the same set of mice. Plots show the mean ± SEM of the protein levels measured in the brains of ten control mice and sixteen *TSC1*+/- mice. Student’s t-test was used to assess statistical significance. Differences were considered statistically significant when p<0.05.

Next, we assessed if the increased tau accumulation observed in 15-month old animals correlated with behavioral abnormalities. The auditory conditioning test measures associative memory, represented by the decrease of the freezing time in response to auditory conditioning (*48*). Compared to *hTau;TSC1*+/+ controls, *hTau;TSC1*+/- mice showed a significant reduction in freezing time after auditory conditioning (**Fig. 6B**), implying that TSC1 insufficiency leads to associative memory impairment. Furthermore, the nesting test was used to estimate executive function of the animals. Nesting ability was measured based on qualitative scoring of amount of torn nestlet material and nest shape (0-7 numerical scale) (*49, 50*). *hTau;TSC1*+/- mice showed decreased nesting scores compared with control mice (**Fig. 6C**), suggesting impaired executive function akin to what is seen in FTD.

We also evaluated whether the loss of TSC1/hamartin protein induced tau acetylation *in vivo*, similar to that observed in *TSC1*+/- neuronal cells. First, we generated an antibody against tau acetyl-lysine 343 (AcK343) (**Fig. S7**) and measured the levels of acetylated tau in mice RSCx lysates. Tau AcK343 levels were strikingly increased in 15-month *hTau;TSC1*+/- animals compared to controls (**Fig. 6D**). Consistent with *TSC1*+/- cells, *hTau;TSC1*+/- mice also demonstrated increased p300 HAT activity and decreased levels of SIRT1 HDAC (**Fig. 6E**). These results confirm that TSC1/hamartin haploinsufficiency leads to acetyl-tau accumulation and behavioral impairments *in vivo* in an age-dependent manner.

## Discussion

In the current study, we validate rare variants in *TSC1* as putative risk factors for tauopathies, demonstrate the accumulation of acetylated tau in *TSC1*+/- neuronal cells and aging mice, and provide a mechanistic basis for tau accumulation in TSC1/hamartin haploinsuffíciency (**Fig. 7**). LOF pathogenic variants in the *TSC1* gene leading to TSC1/hamartin haploinsuffíciency have previously been associated with the juvenile-onset lysosomal storage disease TSC (*12, 51, 52*). In earlier clinical-translational studies, we have shown a linkage between *TSC1* gene and age-related tauopathies (*11, 15*). With this study, we establish the mechanistic basis for this linkage, adding *TSC1* to a set of genes including glucocerebrosidase A (*GBA*), progranulin (*PGRN*) and cathepsin D (*CTSD*) in which pathogenic variants are associated with both childhood developmental and age-associated neurodegenerative diseases (*7, 8, 53*). *GBA, PGRN*, and *CTSD* genes encode for lysosomal resident proteins and therefore pathogenic variants in these genes lead to disrupted autophagy and lysosome storage disorders (*54*). In contrast, *TSC1* haploinsufficent cells exhibit compensated autophagic flux and prevent tau degradation via aberrant post-translational modifications. Therefore, variants in *TSC1* seemingly represent a related but distinct mechanism for inducing neurodegeneration.

**Figure 7.**
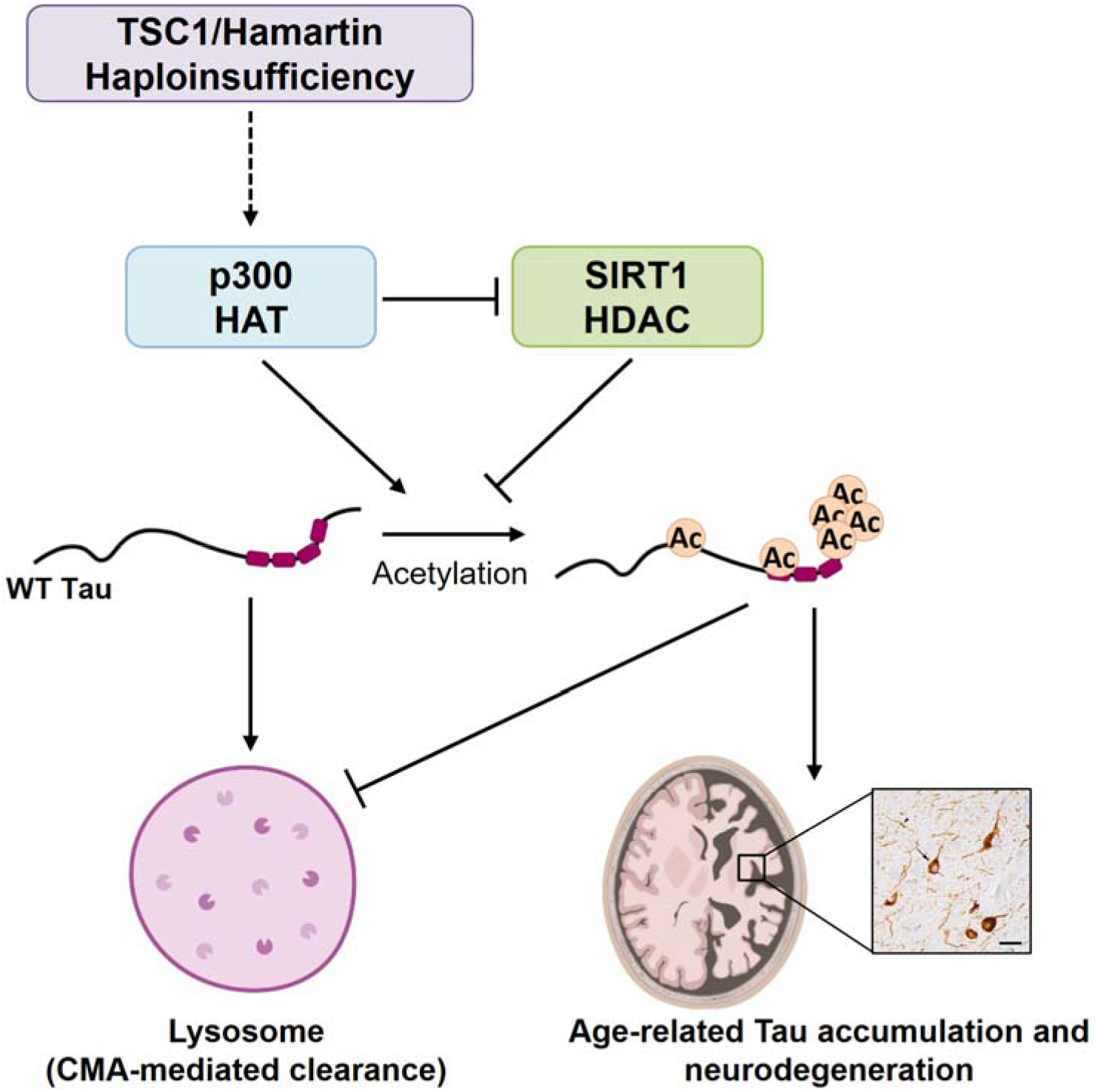
Proposed model for TSC1/hamartin haploinsufficiency impairing tau degradation and increasing risk for tauopathy. Decreased TSC1/hamartin levels induce the activation of p300 HAT and decrease SIRT1 HDAC expression. As a consequence, tau is specifically acetylated at six lysine residues. Acetylated tau fails to be degraded in the lysosomes through chaperone mediated autophagy (CMA) and thus tau accumulates, leading to increased risk of neurodegeneration.

Post-translational modifications on tau can dramatically impact its function, localization and degradation. In this study, we demonstrate that *TSC1* insufficiency leads to specific tau acetylation and thereby reduces its association with lysosomes, specifically those active for chaperone-mediated autophagy. While abnormal tau acetylation has been previously described in tauopathies (*3, 38, 39*), phosphorylation has historically been considered to be the trigger of tau accumulation and aggregation. In contrast, our study implicates acetylation as the key modification leading to tau accumulation through the blockage of its degradation in the lysosomes.

Why tau specifically accumulates in the affected neurons of tauopathy patients remains unknown. Here, we demonstrate that TSC1/hamartin insufficiency promotes p300 HAT activity and decreases SIRT1 HDAC levels. The dysregulation of these enzymes seems to be associated with specific tau acetylation and accumulation in cellular and animal models of TSC1 haploinsufficiency. It has been described that mTOR interacts with p300 and regulates its activity (*55*). Therefore, the function of TSC1 as key protein regulating the mTOR pathway (*21*) could be related with the increased p300 HAT activity observed in TSC1 insufficiency. Interestingly, a new function of TSC1/hamartin as co-chaperone for Hsp90 has been recently reported (*56*). An involvement of Hsp90 in CMA has been previously described (*31*). Although Hsp90 is not required for targeting of CMA substrates to lysosomes, a fraction of Hsp90 bound at the cytosolic side of the lysosomal membrane contributes to stabilization of binding of CMA substrates once they reach the lysosomal membrane (*57*). Therefore, through its role as cochaperone for Hsp90, TSC1/hamartin could potentially regulate tau degradation via CMA. These two functions of TSC1, as a HAT activity modulator and as a co-chaperone, could be essential for the specific accumulation of acetylated tau under TSC1 haploinsufficiency conditions.

Tauopathies are one of the most frequent causes of age-related morbidity, becoming a healthcare challenge and caregivers’ burden; however, no disease-modifying treatments have been generated yet (*58*). To develop effective therapies for this diverse group of disorders, the many different etiologies and pathogenic mechanisms must be better understood and appreciated. The lack of models that recapitulate wild-type, rather than mutant, tau dysfunction has limited our understanding of the molecular mechanisms behind tau accumulation. This distinction is key, as mutant forms of tau utilize alternative pathways for clearance as compared to non-mutant tau (*25, 59*). Thus, models that accumulate non-mutant tau are needed. In this regard, we have developed a powerful genetic model of tauopathy in which human non-mutant human tau accumulates. Although other models of TSC1/hamartin haploinsufficiency were previously generated, most of them have been used to study neuronal development and TSC (*60–64*), but not tau-related neurodegenerative diseases.

Cellular and animal models of TSC1/hamartin haploinsufficiency show increased accumulation of non-mutant tau protein. However, our results do not clarify how exactly the loss of TSC1 leads to the specific accumulation of tau rather than TDP-43 or other neurodegenerative disease proteins (*11*). Although the unanswered question of specificity represents a limitation of this study, the selective neuronal vulnerability to different neurodegenerative disease proteins remains unknown throughout the field. TSC1 insufficient cell lines and mice may provide new models in which to address this highly important and compelling issue.

Here, we have established a mechanistic link between TSC1/hamartin protein, tau acetylation and tau accumulation. Blocking acetylation with a specific p300 HAT inhibitor or promoting deacetylation with a SIRT1 HDAC activator normalizes tau levels in TSC1 insufficiency conditions. Other preclinical studies have shown that the dysregulation of both types of enzymes is involved in tau acetylation and neurodegeneration (*3, 4, 65–68*). Additionally, several clinical trials are currently investigating the neuroprotective effect of the pharmacological modulation of p300 and SIRT1 (*69, 70*). This study supports the translation of p300 and SIRT1 into targets for drug development for the treatment of tauopathies. Furthermore, the newly identified tau acetyllysine residues could represent novel biomarkers for tauopathy diagnosis and progression. Finally, the novel cellular and animal models of non-mutant tau accumulation that we have generated could be used as preclinical models to develop novel disease-modifying treatments, potentially facilitating the successful translation of future therapies to humans.

## Methods

### Human genetics

Two cohorts of tauopathy patients (EOAD; N=228 and PSP; N=501) were analyzed for the presence of rare *TSC1* variants. *TSC1* variant calls were verified via Sanger Sequencing. The allele frequency of rare *TSC1* variants in EOAD and PSP cohort was compared with the allele frequency of these variants in non-Finnish European controls from the Genome Aggregation Database (gnomAD v2.1.1; https://gnomad.broadinstitute.org/variant/9-135771987-C-CGCT?dataset=gnomad_r2_1) and the ADSP Discovery Case Control database (http://www.niagads.org/sites/all/public_files/ADSP%20%20SUMMARY%20PLAN%20revised%20fnl%2041513.pdf), respectively. All participants or their surrogates provided written informed consent to participate in this study and the institutional review boards at each site approved all aspects of the study. A detailed explanation of the methodology used to study the enrichment of *TSC1* variants in tauopathy cases is included as supplementary material.

### Culture and generation of stable SH-SY5Y cell lines

SH-SY5Y neuroblastoma cells (ATCC, #CRL-2266) were cultured in EMEM:F12 medium supplemented with 10% (v/v) heat-inactivated fetal bovine serum (FBS) and 1% penicillin/streptomycin (P/S). *TSC1* heterozygous SH-SY5Y cell lines (*TSC1*+/-) were generated using CRISPR/Cas9 system as previously described (*11*). *TSC2* knockdown (KD) stable cell lines were generated by infecting SH-SY5Y cells with tuberin shRNA (h) lentiviral particles (Santa Cruz Biotechnology, #sc-36762-V) followed by puromycin selection (1μg/mL). To generate stable SH-SY5Y cell lines overexpressing TSC1/hamartin (WT and mutant) and 2N4R tau (WT, acetyl-mimetic K→Q and non-acetylatable K→R), we cloned WT and mutant forms of the *TSC1* and *MAPT/tau* genes in the pLenti-CMV plasmid (Addgene plasmid # 17392) (*71*). Lentiviral particles were generated in HEK293FT cells and used to transduce SH-SY5Y cells as previously described (*11*). Cells expressing the plasmids were selected with neomycin (2μg/uL), expanded and banked. Overexpressed proteins were tagged with myc (N-terminal) and FLAG (C-terminal) tags. Efficiency of protein overexpression was assessed by immunoblotting with antibodies against the myc and FLAG tags.

### Generation and culture of the fibroblast lines

Epithelial fibroblast cells were obtained from the Memory and Aging Center Fibroblast Bank at the University of California, San Francisco. The skin biopsy was performed after obtaining the donor’s written informed consent. The consent allowed for use of tissue by all parties, commercial and academic, for the purposes of research but not for use in human therapy. The informed consent and the protocol of our study was approved by the UCSF Institutional Review Board and Ethics Committee. The biopsy tissue was placed into a tissue culture dish with a coverslip on top. Fibroblasts were fed every 2-3 days with DMEM (ThermoFisher Scientific, #11995073) media supplemented with 10% (v/v) heat-inactivated FBS and 1% P/S. Cell were expanded, tested negative for mycoplasma, banked and used in further experiments.

### iPSC lines generation, genome engineering and characterization

The TSC1-family iPSC lines were generated from epithelial fibroblasts from each member of the family, including three *TSC1* mutation carriers and one non-mutation carrier that was used as a control (**Table S1**). For additional control lines, we used two well-characterized WT iPSC lines: the F11350 iPSC-line obtained from the Karch lab at Washington University School of Medicine (*20*) and PGP-1 iPSC line (GM23338) derived from a 55 year old male obtained from the Coriell Institute for Medical Research (**Table S1**).

Two monoclonal *TSC1*+/- iPSC lines were generated by the company Synthego in the background of the WT PGP-1 iPSC using the CRISPRs/Cas9 gene editing system. Briefly, a sgRNA (5’-ACCAAGGUGUUUACAAGCAU-3’) targeting the *TSC1 gene* was designed and validated before the nucleofection of iPSC, as previously described (*72–74*). To select *TSC1*+/- iPSC clones, genomic DNA was extracted with the QuickExtract kit (Epicentre, #QE09050). PCR was performed using Q5 Hot Start High-Fidelity 2X Master Mix, using 5’-GCAGAACTGTAATGCTGCACAA-3’ and 5’-TCAAGAATCATGGGTCCTACAAAGT-3’ primers. The PCR program was 98°C for 30 s, 30 cycles of 98°C for 10 s, 66°C for 30 s, and 72°C for 25 s, and a final stage at 72°C for 10 minutes. Two *TSC1* heterozygous (*TSC1*+/-) CRISPR lines (B02 and C10) were identified, expanded and banked.

After the generation, both TSC1-family and CRISPR-engineered iPSC lines were verified for the expression of pluripotency markers (SSEA4, Oct4 and SOX2) by immunofluorescence as previously described (*20*) (**Fig. S1-2**). Additionally, chromosomal abnormalities in all iPSC lines were assessed by G-band karyotyping using standard cytogenetic procedures (**Fig. S1-2**).

All iPSC lines were maintained in matrigel (Corning, #354277) coated plates using mTSER media (StemCell Technologies, #05850). For passaging, iPSCs were detached using Accutase (ThermoFisher Scientific, #A1110501) and seeded using mTSER media supplemented with 10μM Rock inhibitor (Y-27632, StemCell Technologies, #72304). Media was replaced every day.

### SH-SY5Y and iPSC differentiation into neurons

For the neuronal differentiation of SH-SY5Y cells, 10,000 cells/cm^2^ cells were seeded in tissue culture plates 24 hours before starting the differentiation through the addition of 10μM of retinoic acid (RA) in EMEM/F12 media supplemented with 10% FBS and 1% P/S. After 6 days of RA treatment, cells were treated for 4 days with 50ng/mL of BDNF in EMEM/F12 media supplemented only with 1% P/S (*74, 75*). Human iPSCs were differentiated into cortical neurons by infection with virus particles containing doxycycline-inducible neurogenin-2 (Ngn2) and puromycin resistance as previously described (*74, 76*). After infection, cells were cultured for 21 days before lysate collection.

### Whole cell lysates and lysosome isolation

For whole lysates, cells were harvested and lysed with Pierce radioimmunoprecipitation (RIPA) buffer (25mM Tris-HCl, pH 7.6, 150mM NaCl, 1% NP-40, 1% sodium deoxycholate, and 0.1% SDS) (ThermoFisher Scientific, #89900) containing protease (MilliporeSigma, #4693124001), phosphatase (MilliporeSigma, #4906837001) and deacetylase inhibitors (3μM trichostatin A (Cayman Chemical Company, #89730) and 10mM niacinamide (MilliporeSigma, #N5535). Then, lysates were centrifuged at 15,000 rpm for 15 min at 4 °C to obtain the supernatant containing soluble proteins.

Lysosomes were isolated from differentiated SH-SY5Y cells following a protocol previously published (*33*). Briefly, 40 million differentiated SH-SY5Y cells were collected and lysed in 0.25M sucrose buffer. After serial centrifugations, a fraction containing both mitochondria and lysosomes was obtained. Lysosomes were isolated and separated from the mitochondrial fraction in a discontinuous density gradient. After centrifugation, lysosomes appear in two aqueous density-dependent layers, the less dense fraction containing CMA+ lysosomes and the denser fraction containing a combinatiom of CMA+ and CMA-lysosomes further separated by differential centrifugation. Lysosomal fractions were resuspended in 0.3M sucrose and centrifuged again at 10,000 × *g* for 5 min. Pellets containing the two lysosome fractions were then diluted in 0.25M sucrose buffer before analysis.

### Generation of anti-tau acetyl lysine 343 antibody, western blotting and protein half-life measurement

Monoclonal anti-tau acetyl lysine 343 (AcK343) antibody was generated in the antibody development facility of Fred Hutchinson Cancer Research Center (Seattle, WA). Mice were immunized using the peptide sequence CVEVKSE(KAc)LDFKD, containing the K343 residue of 2N4R tau. Site specificity of AcK343 was confirmed using acetylated and non-acetylated forms of tau peptides bind to BSA protein (**Fig. S10**).

For western blotting, equivalent amounts of protein were separated on a 4-12% Bis-Tris gel (ThermoFisher Scientific). Immunoblotting was performed as previously described (*74*) using the antibodies listed in supplementary material (Table S2). Immunoreactive bands were visualized using the LI-COR Odyssey CLx imaging system and quantified using the ImageJ software.

To measure the half-life of WT and mutant TSC1/hamartin protein, SH-SY5Y cells expressing FLAG-tagged WT or mutant TSC1/hamartin were seeded into 6-well plates at 1×10^6^ cells/well. Twenty-four hours after seeding, cells were treated with 50μg/mL of cycloheximide (CHX) and lysates were collected at 0, 8, 24, 48 and 72 hours into cold RIPA buffer. Degradation of FLAG-tagged TSC1/hamartin protein was evaluated by immunoblot with an anti-FLAG antibody. Protein half-life was calculated with the following equation: Half-life (T_1/2_)=ln(2)/K, where K=degradation rate constant (hour^-1^).

### Immunoprecipitation and mass spectrometry

Endogenous tau was immunoprecipitated from whole lysates of differentiated SH-SY5Y cells (1mg) using 10μg of the HT7 tau antibody (ThermoFisher Scientific, #MN1000) and the crosslink magnetic immunoprecipitation kit (Thermo-Pierce, # PI88805) following the manufacturer’s instructions. Immunoprecipitated proteins were separated on a 4–12% Bis-Tris gel and stained with SafeStain (ThermoFisher Scientific, #LC6060). A broad band around 50 kDa was cut out from the gel, digested with trypsin and analyzed by mass spectrometry. The identification of peptides and post-translational modifications in tau-enriched samples was performed at the UCSF Mass Spectrometry Facility following standard protocols (*77, 78*).

### Quantitative PCR analysis

SH-SY5Y cells were differentiated in 10 cm plates and either collected right after differentiation or treated with A485 (15μM) or DMSO for 72 hours prior to collection. mRNA was extracted using the RNeasy kit (Qiagen, #217004) per the manufacturer’s protocol. Extracted mRNA was converted to cDNA by PCR using SuperScript III Reverse Transcriptase (Invitrogen, #18080-044). The expression of the *MAPT*/tau gene and *SIRT1* gene was measured by qPCR as previously described (*79*). *GAPDH* and *actin* were used as controls.

### Autophagy analysis

Measurement of macroautophagy in differentiated SH-SY5Y cells was performed as previously described (*29*). Briefly, cells were treated with the V-ATPase inhibitor bafilomycin A1 (100nM, BafA1) for 6 hours to block protein degradation in the lysosomes, lysates were collected, and then LC3-II protein levels were assessed by immunoblot. An increase in LC3-II levels after BafA1 treatment corresponds with an increase in autophagy flux.

Macroautophagy activity in fibroblasts was measured after transduction with lentivirus carrying an mCherry-GFP-LC3 tandem construct (*27*). Fibroblasts were plated on glass-bottom 96 well plates, fixed with 4% paraformaldehyde (PFA), and fluorescence was read in both red and green channels using high-content microscopy (Operetta, Perkin Elmer). Images of nine different fields per well were captured, resulting in an average of 2,500-3,000 cells per view. The number of particles/puncta per cell was quantified using the “particle identifier” function in the cytosolic region (*80*). Values were presented as number of puncta per cell section. Positive puncta for both fluorophores correspond to autophagosomes, whereas those only positive for the red fluorophore correspond to autolysosomes. Autophagic flux was determined as the conversion of autophagosomes (yellow) to autolysosomes (red only puncta) (*27*)

### Immunofluorescence, confocal laser scanning microscopy and colocalization analysis

Undifferentiated and differentiated SH-SY5Y cells were seeded in microscopy chambers (ThermoFisher Scientific, #155382). When confluent, cells were fixed for 30 min in 4 % PFA in PBS. Then cells were washed twice with 2mg/mL glycine and blocked with 2% BSA in PBS for 1 hour. Cells were permeabilized with 0.1% saponin, incubated with primary antibodies for 1 hour at room temperature, and then incubated with the corresponding secondary antibodies for 45 min at room temperature. ProLong Gold Antifade Reagent with DAPI was used for nuclear staining (Cell Signaling Technology, #8961S). Stains were visualized in the High Speed Widefield confocal (UCSF Nikon core) using a water immersion 63x objective using green FITC-488 excitation and far-red 647 excitation. Pearson’s correlation coefficient measured with Fiji/Image J was used to estimate the colocalization between both channels.

### Measurement of p300 histone acetyltransferase activity

The acetyl transferase activity of p300 was measured using the HAT activity colorimetric assay kit (MilliporeSigma, #EPI001) per manufacturer’s instructions. Briefly, differentiated SH-SY5Y cells were collected, washed with PBS, and proteins were extracted with CHAPS buffer (50mM HEPES, pH 7.4, 40mM NaCl, 2mM EDTA, 1mM orthovanadate, 50mM NaF, 10mM pyrophosphate, 0.3% CHAPS) including protease and phosphatase inhibitors. Then, 75μg of whole lysate was incubated with the kit assay mix and run in triplicate in 96 well plates for 1 hour at 37°C. Values were read at O.D. 440nm using the Tecan M200 Pro plate reader. To block the activity of p300 HAT, lysates were treated for 30 min at 37°C with the specific p300 inhibitor, A485. The activity of p300 HAT was assessed by subtracting the remaining HAT activity after A485 treatment from the initial values in non-treated lysates.

### Transgenic mouse generation and brain lysate extraction

*hTau;TSC1*^Syn1+/+^ and *hTau;TSC1*^Syn1+/-^ transgenic mice were bred and maintained in-house at Washington University in St. Louis School of Medicine. hTau mice hemizygous (Cre+ or *hTau;TSC1*^Syn1+/-^) or homozygous (Cre- or *hTau;TSC1*^Syn1+/+^) for TSC1 were age-matched and randomized across both sex and litter for use in studies at either 6 months old (n=3 *hTau;TSC1*^Syn1+/+^ and n=6 *hTau;TSC1*^Syn1+/-^) or 15 months old (n=10 *hTau;TSC1*^Syn1+/+^ and n=16 *hTau;TSC1*^Syn1+/-^). Methodology for the generation of transgenic mice, behavior analysis, euthanasia, tissue processing, immunohistochemistry, imaging and quantification is included as supplementary material.

For protein quantification studies, the retrosplenial cortex (RSCx) was isolated and lysed by mechanical disruption in cold RIPA buffer. To measure tau acetylation, the heat stable protein fraction, containing tau protein, was extracted (*81*) by centrifugation at 15,000 rpm at 4°C for 15 min. 20uL of the resulting supernatant containing soluble proteins was heated at 95 °C for 30 min. After the heat shock, samples were centrifuged again at 15,0000 rpm at 4°C for 20 min. The supernatant corresponding to the heat stable fraction was collected and analyzed by immunoblot.

### Statistical analysis

Student’s t-test, one-way and two-way ANOVA statistical analyses were performed using GraphPad Prism 6. Bonferroni’s analysis was performed to analyze the statistical significance between multiple groups. Plots show means ± standard error (SEM) and p-values for all the experiments performed.

## Acknowledgments

We gratefully acknowledge all the members of the Kao lab for the critical review of the manuscript. Additionally, we thank Dr. Matt Jacobson, Dr. David Agard and Dr. William Seeley for their thoughtful advice.

## Author contributions

CA and AWK conceived the project. CA performed the molecular biology experiments in cellular models. KMS and TMM designed and performed the experiments in mouse models. ARA generated and banked the patient’s fibroblasts lines. EGG, JSY, EEM, BD, GDS and EMR performed genetic analysis. AS and AMC measured autophagy in fibroblasts. KL and ALB performed the mass-spectrometry experiments and analysis. CA and AWK wrote the paper with input from many authors.

## Data and materials availability

The authors confirm that the data supporting the findings of this study are available within the article and/or its supplementary materials.

