## supplementay methods and figures for "*TSC1* loss-of-function increases risk for tauopathy by inducing tau acetylation and preventing autophagy-mediated tau clearance"

### Supplementary Materials and Methods

#### *Analysis of TSC1 variants in EOAD*

The cohort of EOAD cases (N=228) analyzed for enrichment of rare *TSC1* variants has been previously described (*1*). Briefly, all participants were clinically assessed at the University of California, San Francisco (UCSF) Memory and Aging Center and were negative for known Mendelian variants associated with neurodegenerative diseases. Genomic DNA isolated from whole blood underwent library preparation for sequencing via Covaris shearing, end repair, adapter ligation, and PCR using standard protocols. Libraries were sequenced at the HudsonAlpha Institute for Biotechnology on the Illumina HiSeq X platform to achieve 30x coverage. Alignment of raw sequencing reads to the hg19 reference genome was done with bwa-0.7.12 (*2*). Indel realignment, base recalibration, and gVCF generation were done with GATK 3.3 (*3*), while GATK 3.8 was used for joint variant calling across all gVCFs. The cohort-wide VCF was quality filtered to retain variants in which ≥ 95% of sites have a minimum GQ of 20 and DP of 10, and a variant level filter of VQSLOD > -3. KING 2.1.2 (*4*) was used to check for cryptic relatedness and exclusion of related individuals (up to 4th degree relatives using IBD segment analysis). All participants in this study were confirmed to be of European ancestry by principal component analysis using plink 1.9 (*5*) compared to 1000 genomes data (*6*) and analysis using ADMIXTURE 1.3.0 (*7*).

#### *Analysis of TSC1 variants in PSP*

Whole exome sequencing (WES) data from a cohort of 775 PSP patients was analyzed for variants in *TSC1* gene. Samples were sequenced initially, and additional sequencing was done when coverage did not reach 20X for more than 80% targeted region and 10X for more than 90% of the targeted regions. PSP samples were enriched for exonic DNA using Nimblegen's VCRome v2.1 (36Mb) capture kit. PSP sequencing data was analyzed using the in-house DNA Resequencing analysis workflow (DRAW) (*8*). First, we mapped sequencing reads to the GRCh37 reference genome using BWA 0.7.5 (*9*). We saved read alignments as BAM files and used PICARD (http://picard.sourceforge.net/) to mark duplicate reads. Additional sequencing was conducted to reach targeted coverage (>85% and >75% of targeted regions reach >20x and

>10x respectively). We merged data from multiple sequencing experiments from the same individual using SAMtools (http://samtools.sourceforge.net) (*10*). GATK was used to perform local read realignment near known insertions or deletions (indels) sites and base quality score recalibration (BQSR) (*11*). Variants were called from the processed BAM files using GATK HaplotypeCaller (*12*). Calls were restricted to the exome capture regions, each expanded with 30bp flanking regions on both sides. A joint genotyping analysis of the all individual gVCFs were performed to produce a project level VCF (pVCF). VerifyBamID (http://genome.sph.umich.edu/wiki/VerifyBamID) was used to check for possible sample contaminations in the PSP sequencing data (freemix) (*13*). Concordance with SNP array data was validated using common SNVs (MAF > 0.05) in the exome capture intervals; samples with discordance rate greater than 2% were removed. Duplicate/related samples in the same SNP panel were identified using PLINK (*5*); for duplicate/related samples (IBD>0.125), the one with the highest call rate was retained and the rest were removed. 637 PSP samples passed the initial sample and sequence quality checks. Of these, 550 PSP samples of European origin (non-Hispanic white) were selected to be included in the ADSP GRCh38 joint-called 20K WES dataset (https://dss.niagads.org/niagads-dss-releases-20k-whole-exomes/).

The initial ADSP 20K WES dataset release is comprised of 19,922 whole exomes called on GRCh38. Nine different studies contributed to the dataset, including 10,088 ADSP Discovery Case Control WES samples, and 550 PSP WES samples (http://www.niagads.org/sites/all/public_files/ADSP%20%20SUMMARY%20PLAN%20revised%20fnl%2041513.pdf). The ADSP Discovery cohort samples were sequenced in three sequencing centers; Broad Institute used the Illumina Rapid Capture Exome (ICE) kit, Baylor and WashU used the Nimblegen's VCRome v2.1 kit. Briefly, the WES reads were mapped to the GRCh38 reference and variants were joint-called using the Genome Center for Alzheimer’s Disease (GCAD) pipeline, VCPA 1.1, a functionally equivalent CCDG/TOPMed pipeline (*14*). Variant and sample quality checks were performed using the ADSP QC pipeline (*15*). More information about the ADSP 20K WES dataset can be found on the NIAGADS Data Sharing Service ng00067 dataset page (https://dss.niagads.org/datasets/ng00067/). WES data for the 550 PSP samples and 4,182 cognitively normal ADSP Discovery control samples of European decent was extracted from the ADSP 20K WES dataset pVCF, which was accessed through the

NIAGADS Data Sharing Service (https://dss.niagads.org/). Subsequent QC and analyses were restricted to variants in the exome capture kit consensus intervals.

We combined GRCh38 genotype calls from the 550 PSP samples and 4,182 ADSP controls with the 1000 Genomes project Phase 3 samples (*16*), and applied principal components analysis (PCA) using independent SNPs with site-missing rate < 0.05, MAF > 0.05, located in both target region of PSP and ADSP, followed by LD pruning (plink --indep-pairwise 50 5 0.2). 540 PSP samples of European descent (non-Hispanic white) were retained after dropping 10 PSP samples as outliers. 4,167 ADSP control samples of European descent were retained after dropping 15 ADSP control samples as outliers. With the remaining PSP and ADSP control samples, we performed a second PCA with only case and control samples to identify additional cohort outliers; 507 PSP samples and 3,852 ADSP control samples were retained. Using SNVs passing ADSP QC pipeline variant-level quality checks, sample-level quality was checked for PSP and ADSP controls separately using the following criteria: (1) number of singleton or doubleton variants (variants with only one or two alternative allele carriers in the entire dataset) < 6 SDs above the population mean. (2) missing call rates < 20%. (3) Inbreeding coefficient < 6 SDs above the population mean. 501 PSP and 3,821 ADSP control samples passed all criteria and were included in the subsequent *TSC1* variant analyses. We queried the PSP – ADSP control WES pVCF for variants in the *TSC1* gene using BCFtools (http://www.htslib.org/doc/bcftools.html). PSP and ADSP control samples with heterozygous or homozygous alternate allele calls for TSC1 variants rs2234980 and rs118203742 were extracted using BCFtools. The *TSC1* WES variant calls were then verified via Sanger Sequencing.

The PSP *TSC1* variant statistics were calculated using the PSP and ADSP controls validated allele counts. Samples with rs118203742 or rs2234980 variant calls were excluded from the statistical calculations if DNA was not available for validation testing, or if the DNA sample failed to amplify, and the variant call could not be validated. Samples with variant calls that validated as homozygous Reference were retained and the corresponding genotype call and allele count were corrected accordingly. Only validated variant calls were included in the rs118203742 and rs2234980 *TSC1* variant statistics. The PSP and ADSP control allele frequencies, Odds Ratio, and One-sided Fisher's Exact Test were calculated using Python packages Pandas (*17*) and SciPy (*18*).

*Sanger Sequencing*

Genomic DNA (~50ng) was amplified using a SimpliAmp Thermal Cycler (Applied Biosystems) in a 20ul reaction volume with HotStarTaq Master Mix (Qiagen) in the presence of 2uM primers (TSC1_rs118203743-F1: TGCCGTCCTCATCACACTG, TSC1_rs118203743-R2: TCCCTCCATATGGCCACAG, IDT). The PCR conditions used were: 95°C 15min followed by 30 cycles of 95°C 20sec, 55°C 30sec, 72°C 2min with a final extension of 72°C 7min. The amplified PCR products were prepared for Sanger sequencing by adding ExoSAP-IT (USB) and incubating at 37°C for 45min followed by 80°C for 15min. The PCR products were then Sanger sequenced using the BigDye® Terminator v3.1 Cycle Sequencing kit (Part No. 4336917 Applied Biosystems). The sequencing reaction contained BigDye® Terminator v3.1 Ready Reaction Mix, 5X Sequencing Buffer, 5M Betaine solution (Part No. B0300 Sigma) and 0.64uM sequencing primer (same as the PCR primers) in a total volume of 5ul. The sequencing reaction was performed in a SimpliAmp Thermal Cycler (Applied Biosystems) using the following program: 96°C 1min followed by 25 cycles of 96°C 10sec, 50°C 5sec, 60°C 1min15sec. The products were cleaned using XTerminator and SAM Solution (Applied Biosystems) with 30min of shaking at 1800rpm followed by centrifugation at 1000 rpm for 2min. The sequencing products were analyzed on a 3130xl Genetic Analyzer (Applied Biosystems) and the sequencing traces were analyzed using Sequencher 5.4 (Gene Code).

*Transgenic mice*

Mice expressing Cre recombinase under the synapsin (Syn1) promoter (Syn1-Cre; 003966 Jackson Laboratories, Bar Harbor, ME) and mice expressing floxed TSC1 alleles (TSC1Fl/Fl; gift from Michael Wong, Washington University in St. Louis) were bred onto a mouse tau knockout (mTau-/-) genotype. The resulting crosses were then bred to mice expressing all six isoforms of human tau (hTau) in the absence of mouse tau (*19*). Mice were genotyped using tail DNA for human tau (Forward: 5’-ACTTTGAACCAGGATGGCTGAGCCC-3’, Reverse: 5’-CTGTGCATGGCTGTCCACTAACCTT-3’), EGFP (indicative of mouse tau knockout; Forward: 5’-TGCTCAGGTAGTGGTTGTCG-3’, Reverse: 5’-TGCTCAGGTAGTGGTTGTCG-3’), TSC1 (Forward: 5’-AGGAGGCCTCTTCTGCTACCACTTTT-3’, Reverse (wild type allele): 5’-GAAGGCAGCTCCGACCATGAAGTGC-3’, Reverse (floxed allele): 5’-

ACGTAGCCGGCTAACGTTAACAACC-3’), and Cre recombinase (Forward: 5’-ACGAACCTGGTCGAAATCAGTGCG-3’,Reverse:5’-CGGTCGATGCAACGAGTGATGAG-3’). hTau mice hemizygous (Cre+ or *hTau;TSC1*$^{Syn1+/-}$) or homozygous (Cre- or *hTau;TSC1*$^{Syn1+/+}$) for *TSC1* were age matched and randomized across both sex and litter for use in studies at either 6 or 15 months of age.

Mice were housed in a double-barrier facility under 12:12 light/dark conditions and given access to food and water *ad libitum*. All husbandry, behavior and other procedures were approved by and performed according to the Washington University in St. Louis Institutional Animal Use and Care Committee.

*Behavior analysis*

At 15 months of age, *hTau;TSC1*$^{Syn1+/+}$ and *hTau;TSC1*$^{Syn1+/-}$ transgenic mice underwent behavioral analysis. For the nesting behavior test, mice were given 3g of untorn nestlet material at the start of their 12-hour dark cycle. Approximately 15 hours later, the resulting nests were evaluated for the amount of nestlet material remaining untorn and for nest quality on a 7-point scale, with a score of 0 being poor and a score of 7 being a near perfect nest (*20*). Nest images were scored by an observer blinded to mouse genotype. The auditory conditioning test, as part of the conditioned fear task, was conducted and analyzed by the Animal Behavior Core at Washington University in St. Louis School of Medicine. Technicians blinded to mouse genotype trained and tested mice using two clear, plastic conditioning chambers per a previously described protocol (*21*). In brief, mice were placed into the training conditioning chamber for 5 min and freezing behavior was quantified using FreezeFrame image analysis software (Actimetrics, Evanston, IL). After a 2 min baseline period, a conditioned stimulus (80 dB tone) was presented for 20 sec followed by an unconditioned stimulus (1 sec, 1.0 mA continuous foot shock). The tone-shock pairing was repeated each minute for a 2 min period. Forty-eight hours later, mice were evaluated for auditory conditioning by placing them into the other conditioning chamber separate from the environment in which the tone-shock occurred. Freezing behavior was quantified during a 2 min baseline period and over a subsequent 8 min period in which the auditory cue (conditioned stimulus) was presented.

*Euthanasia and tissue processing*

At either 6- or 15-months of age, mice were euthanized by transcardial perfusion with ice-cold 1X phosphate buffered saline (PBS). Brain tissue was removed, and the left and right hemispheres were separated for histological and biochemical analyses, respectively. The left hemisphere was dropped into cold 4% PFA, fixed overnight, and then transferred to 30% sucrose in 1X PBS for 24-48 hours. The tissue was then frozen in cold (-25°C to -35°C) isopentanes and stored at -80°C until use. The right hemisphere was sectioned into regions and flash frozen in liquid nitrogen. Tissue was stored at -80°C until use.

*Immunohistochemistry in mouse brains*

Frozen left brain tissue was sectioned at 40µm using a sliding microtome and stored free-floating in a cryoprotectant solution (30% ethylene glycol, 30mM phosphate buffer, 15% w/v sucrose, in $H_2O$). For immunohistochemistry of phosphorylated tau, sections (n=6-7 per brain) were washed in 1X Tris-buffered saline (TBS) and incubated in 3% hydrogen peroxide in 0.25% Triton X-100/TBS to quench endogenous peroxidases for 30 min. Sections were then blocked in 5% nonfat dry milk in 1X TBS for 1 hour before incubating in primary antibody (PHF1, 1:1000; generous gift from Dr. Peter Davies) overnight at 4°C. The next day, sections were washed in 0.05% Triton X-100/TBS and incubated in secondary antibody (goat anti-mouse IgG1-biotin, 1:2000; Bio-Rad, Hercules, CA) diluted in 20% Superblock™ Blocking Buffer (ThermoFisher Scientific, #37580) in 0.05% Triton X-100/TBS for 2 hours. Sections were washed in 0.05% Triton X-100/TBS before a 1-hour incubation in avidin/biotin complex using the Vectastain Elite ABC-HRP kit, (1:400; Vector Laboratories, Burlingame, CA, #PK-6100) diluted in 20% Superblock™/0.05% Triton X-100/TBS. Finally, sections were washed (0.05% Triton X-100/TBS) and developed with 3,3'-diaminobenzidine (DAB) peroxidase substrate kit (Vector Laboratories, #SK-4100). Sections were washed in 1X TBS and mounted onto glass slides to dry overnight. The following day, slides were dehydrated in an ethanol gradient and coverslipped with Cytoseal XYL (ThermoFisher Scientific, #8312-4).

*Imaging and quantification*

Slides were imaged at 20X magnification under bright-field settings on a NanoZoomer 2.0-HT whole slide imaging system (Hamamatsu Photonics K.K., Hamamatsu City, Japan). Automatically captured images were acquired using NDP.scan 2.5 software and viewed/exported using NDP.view. *hTau;TSC1* histology imaging was supported by the Hope Center Alafi Neuroimaging Lab and a P30 Neuroscience Blueprint Interdisciplinary Center Core award to Washington University in St. Louis (P30 NS057105). Sections were blinded to genotype and analyzed for the percent area of phosphorylated tau (PHF-1) reactivity within the retrosplenial cortex using ImageJ (NIH).

**Supplementary Figures**

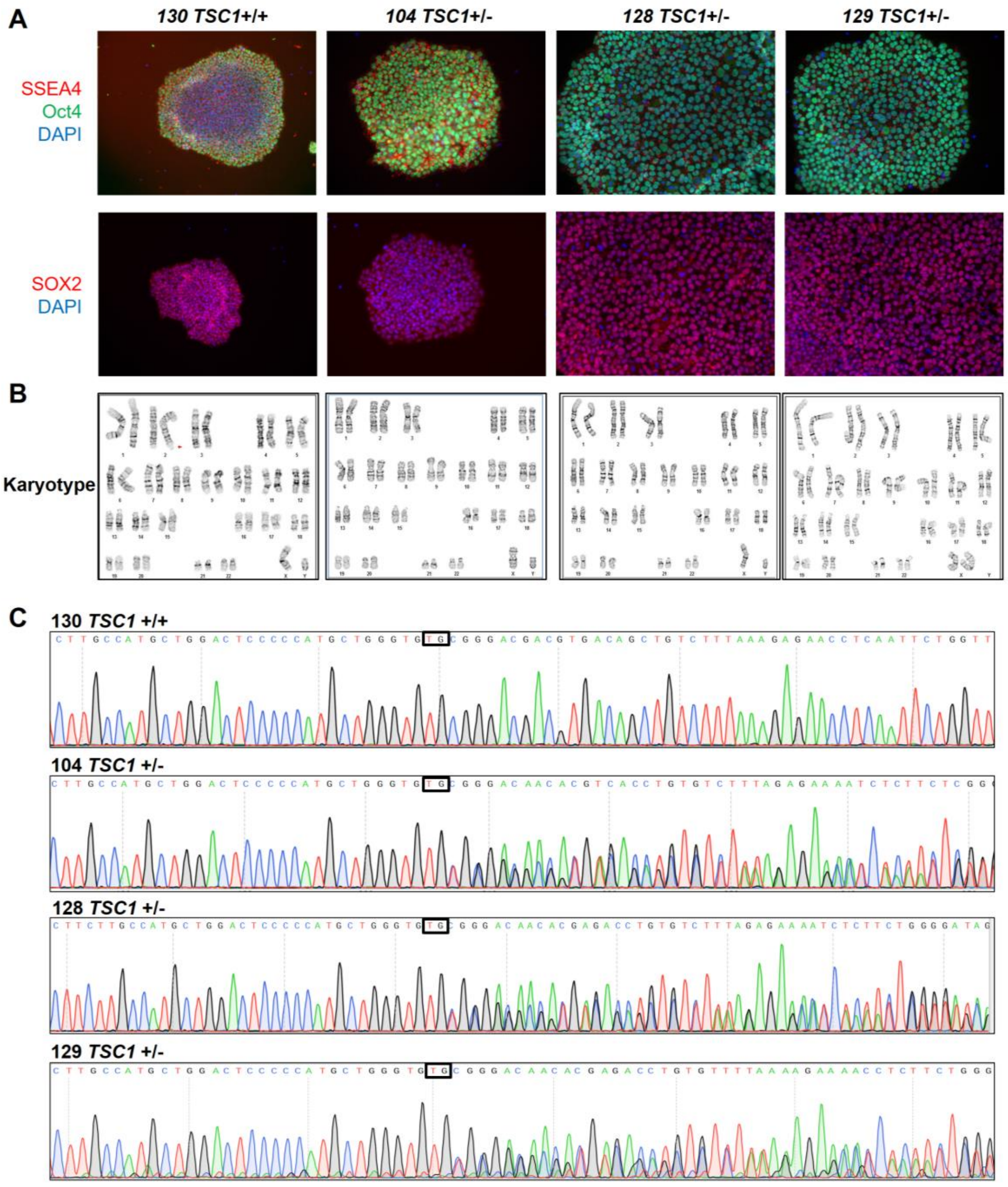


**Supplementary Figure 1. Characterization of iPSC lines from *TSC1* mutation family members.** **A**) Immunofluorescence showing positive staining of the pluripotence markers SSEA4 (red), Oct4 (green) and SOX2 (purple). Nuclei were stained with DAPI (blue). **B**) G-band karyotyping showing that all the iPSC lines used display no chromosomal abnormalities. **C**) Sanger sequencing confirming the presence or absence of the c.62_63inTG mutation in the *TSC1* gene.

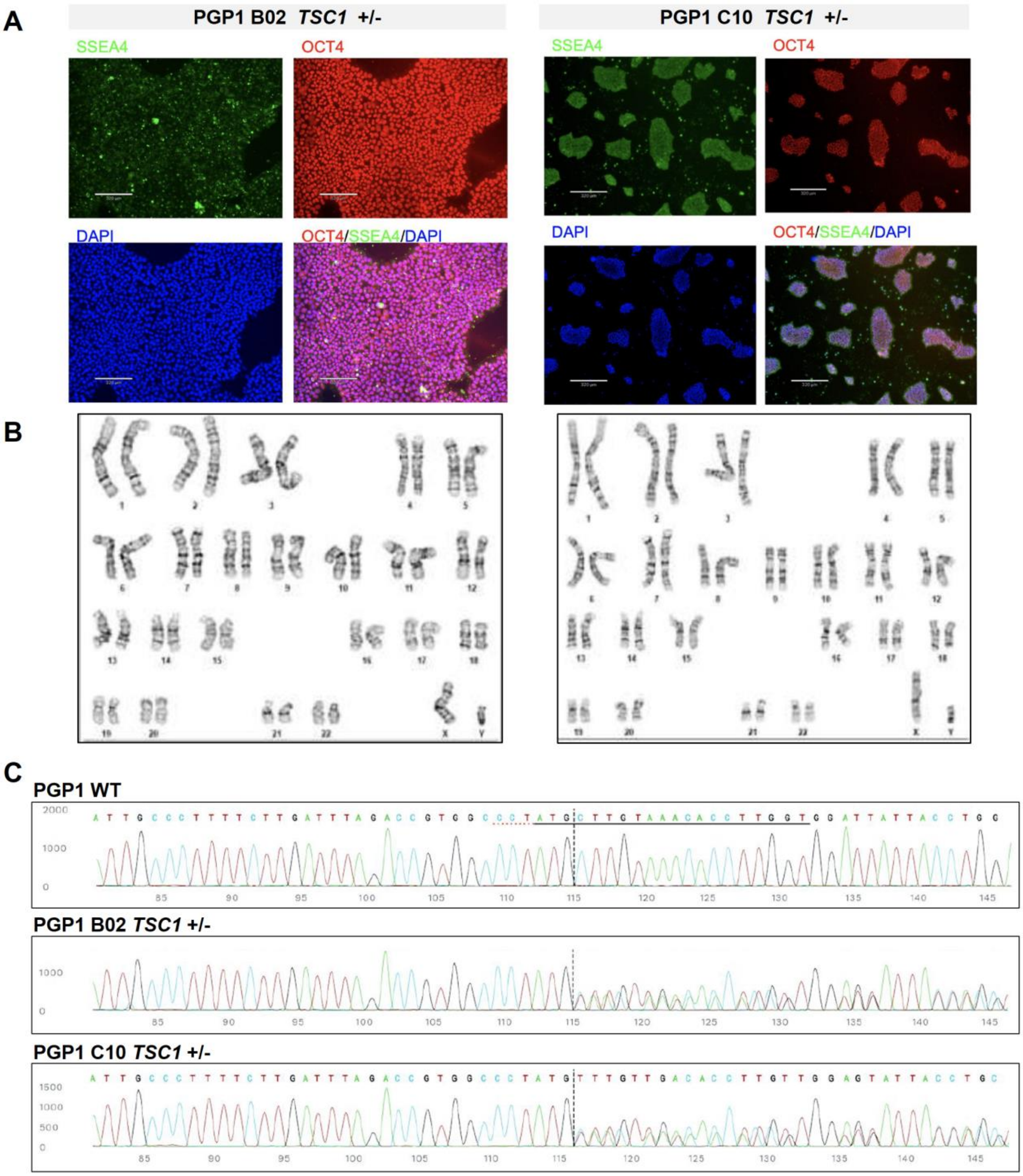


**Supplementary Figure 2. Characterization of CRISPR/Cas9 engineered PGP1 iPSC lines.** **A**) Immunofluorescence showing positive staining of the pluripotence markers SSEA4 (green) and Oct4 (red) in two independent isogenic *TSC1*+/- PSP1 clones. Nuclei were stained with DAPI (blue). **B**) G-band karyotyping showing that all the iPSC lines used display no chromosomal abnormalities. **C**) Sanger sequencing confirming the CRISPR edition in heterozygosis.

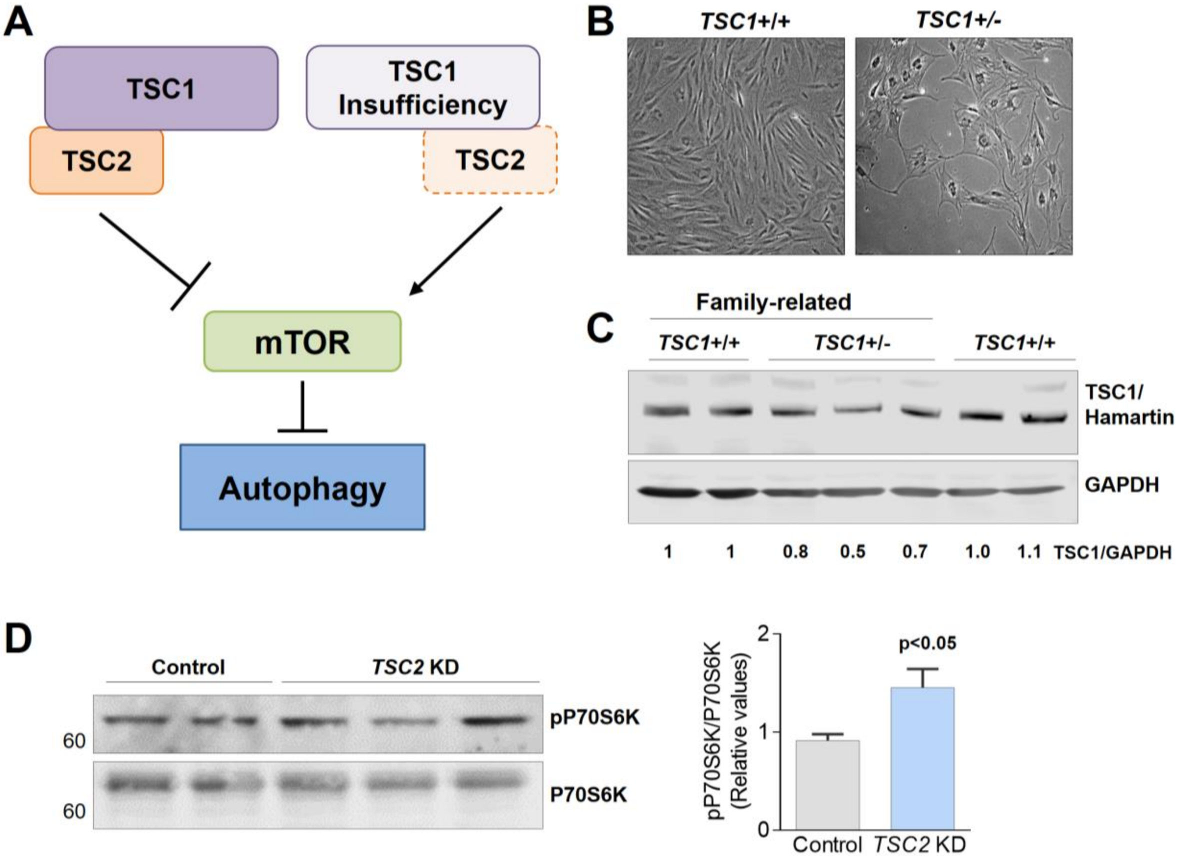


**Supplementary Figure 3. A)** Diagram representing the effect of TSC1/hamartin protein insufficiency in the mTOR pathway and autophagy. **B)** Representative images of wild-type fibroblasts (*TSC1*+/+) and fibroblasts carrying the c.62_63insTG variant in *TSC1 (TSC1*+/-). **C**) Immunoblot showing the TSC1/hamartin levels in *TSC1*+/+ and *TSC1*+/- fibroblasts used in this study. **D)** Immunoblot showing the levels of phosphorylated and non-phosphorylated P70S6K protein control and *TSC2* knockdown (KD) SH-SY5Y cells. The increased phosphorylation of P70S6K protein reflects the overactivation of mTOR pathway. Plot shows the mean ± SEM of four independent experiments performed with all the cell lines used in this study.

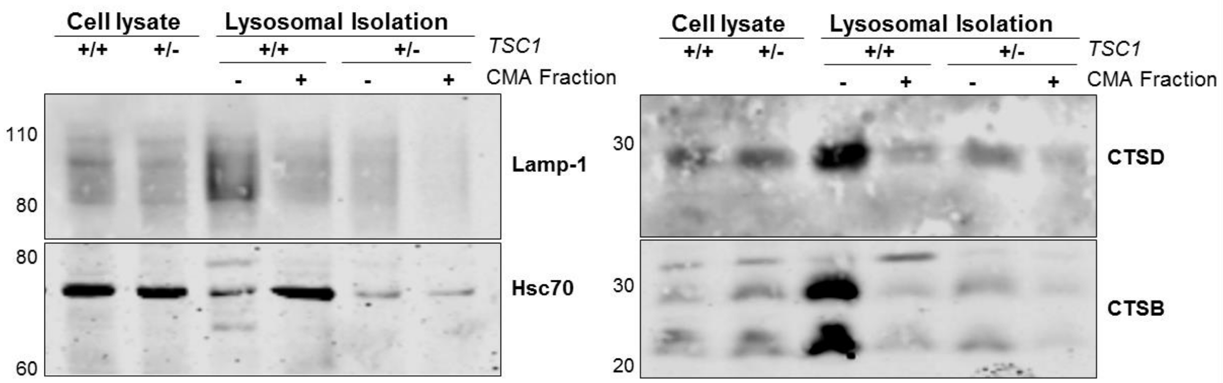


**Supplementary Figure 4. Validation of the lysosome fractionation in differentiated SH-SY5Y cells.** Whole cell lysate and CMA+ and CMA- lysosome fractions were stained with well-established lysosomal markers such as the lysosomal membrane proteins Lamp-1, the CMA chaperone Hsc70, and the lysosomal proteases cathepsin (CTS) D and CTSB. These immunoblots demonstrate the effectiveness of the lysosome isolation procedure.

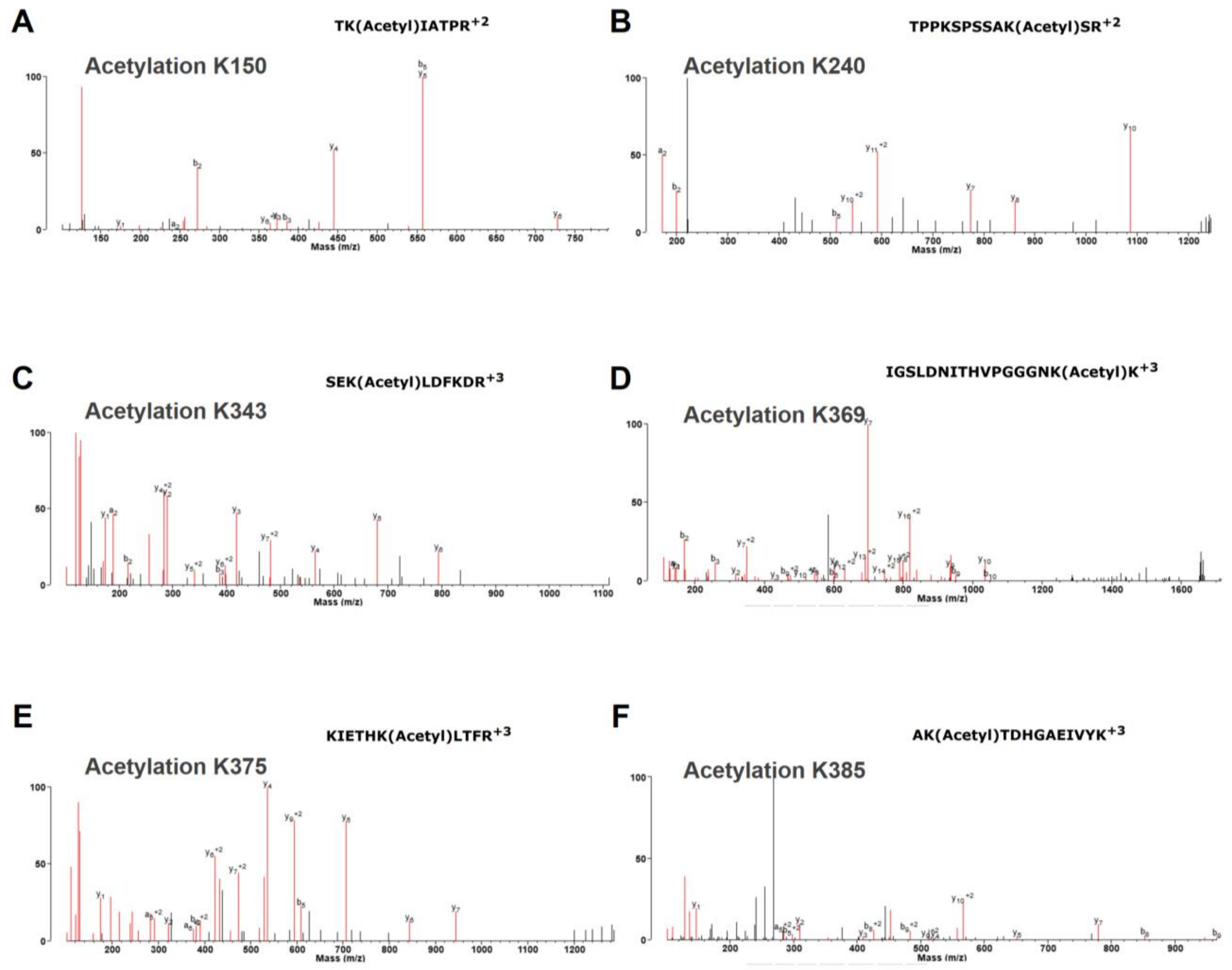


**Supplementary Figure 5.** Mass spectrometry analysis confirming the presence of lysine acetylation in differentiated SH-SY5Y cells. Tau was immunoprecipitated with the HT7 antibody, separated by SDS-PAGE followed by gel excision. The mass spectrometry analysis identified 6 different acetylated peptides (AcK150, AcK240, AcK343, AcK369, AcK375 and AcK385) in *TSC1*+/- differentiated SH-SY5Y cells. The corresponding m/z spectrums for each acetylation site are shown. Complete information about PTMs identified in this study, including acetylation and phosphorylation, is presented in the following links: link1 and link2.

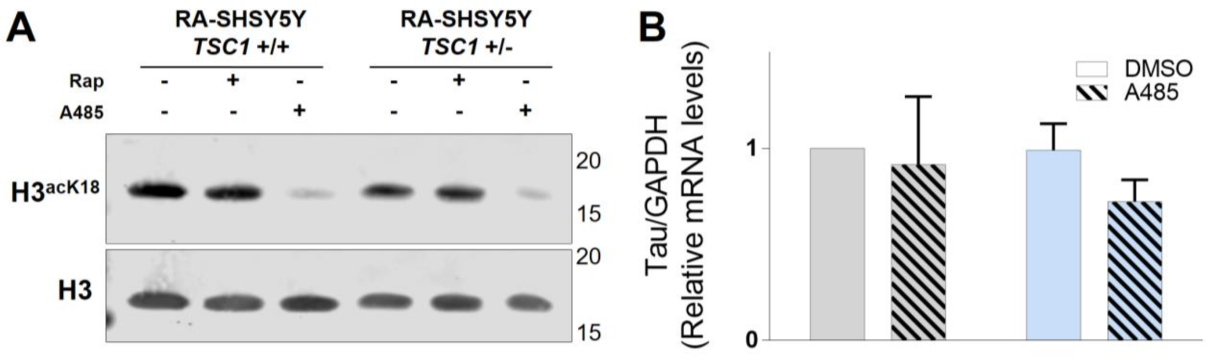


**Supplementary Figure 6. A)** Representative immunoblot showing that differentiated SH-SY5Y cells treated during 72 hours with the p300 inhibitor A485 (15µM) displayed decreased $H3^{aK18}$ acetylation. These data confirm that A485 treatment was effective at decreasing p300 activity. **B**) Plot showing tau mRNA levels from untreated and A485 treated SH-SY5Y cells demonstrating that the inhibition of p300 HAT activity does not affect tau expression. Experiments were performed in triplicates. Plot shows the mean ± SEM.

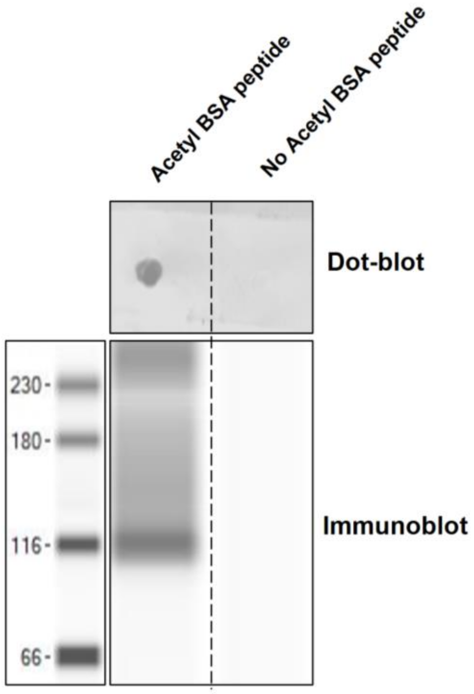


**Supplementary Figure 7.** Validation of the in-house made AcK343 monoclonal antibody. Acetyl-K343 and non-acetyl-K343 BSA peptides were incubated with 1:10 dilution of the antibody overnight. Positive staining only with the Acetyl-K343 BSA peptide by dot blot and immunoblot.

| Line Name | Age | Gender | Familiar history | *TSC1 +/-* | CRISPR engineered? |
|---|---|---|---|---|---|
| iPSC MHF 104 | 55 | Male | Proband | Yes | No |
| iPSC MHF 128 | 53 | Male | Sibling | Yes | No |
| iPSC MHF 129 | 44 | Female | Half sibling | Yes | No |
| iPSC MHF 130 | 70 | Male | Half sibling | No | No |
| iPSC F11350 | Unknown | Male | None | No | No |
| PGP1 WT | 55 | Male | None | No | No |
| PGP1 TSC1 +/- B02 | 55 | Male CRISPR | None | Yes | Yes |
| PGP1 TSC1 +/- C10 | 55 | Male CRISPR | None | Yes | Yes |

**Supplementary table 1**. Demographic characteristics of the iPSC lines used in this study. Highlighted in red are the lines showing TSC1/hamartin haploinsufficiency. iPSC lines in black represent the WT iPSC lines used as controls.

| Antibody | Host | Company (catalog number) | Use and dilution |
|---|---|---|---|
| **M2 FLAG** | Mouse | Sigma (F1804) | Western blot (1:500)/ Immunostaining (1:50) |
| **Hamartin** | Rabbit | Abcam (ab40872) | Western blot (1:500) |
| **Phospho P70S6$^{Thr389}$ Kinase** | Rabbit | Cell Signaling (9234) | Western blot (1:500) |
| **P70S6 Kinase** | Rabbit | Cell Signaling (49D7) | Western blot (1:1000) |
| **Total tau (HT7)** | Mouse | Thermo Scientific (MN1000) | Western blot (1:500)/ Immunostaining (1:50) |
| **Phospho tau$^{Ser396,Ser404}$ (PHF1)** | Mouse | Dr. Peter Davies lab | Western blot (1:500) |
| **GAPDH** | Rabbit | Abcam (ab8245) | Western blot (1:1000) |
| **LC3B** | Rabbit | Sigma (L7543) | Western blot (1:250) |
| **Alpha/beta tubulin** | Rabbit | Cell Signaling (2148) | Western blot (1:1000) |
| **TSC2/tuberin** | Rabbit | Cell Signaling (4308) | Western blot (1:500) |
| **Ac-K343 tau** | Mouse | In house monoclonal antibody (mice serum) | Western blot (1:20) |
| **Lamp2-Alexa 647 (H4B4)** | Mouse | Biolegend (CD107b) | Immunostaining (1:100) |
| **LAMP1 XP** | Rabbit | Cell Signaling (D2D11) | Western blot (1:500) |
| **Hsc70** | Mouse | Abcam (ab2788) | Western blot (1:1000) |
| **CTSD** | Rabbit | ThermoFisher (PA5-72181) | Western blot (1:1000) |
| **CTSB** | Rabbit | ThermoFisher (PA5-14255) | Western blot (1:1000) |
| **H3$^{acK18}$** | Rabbit | Abcam (ac1191) | Western blot (1:250) |
| **H3** | Mouse | Cell Signaling (3638) | Western blot (1:500) |
| **SIRT1** | Mouse | Cell Signaling (8469) | Western blot (1:500) |
| **TDP-43** | Rabbit | Proteintech (12892-1-AP) | Western blot (1:1000) |

**Supplementary table 2.** List of the antibodies used in this study.
